# The tumor-maintaining function of UTX/KDM6A in DNA replication and the PARP1-dependent repair pathway

**DOI:** 10.1101/2024.05.31.596824

**Authors:** Lin-Wen Yeh, Je-Wei Chen, Jia-Yun Yeh, Mei-Han Kao, Hsiao-Chin Hong, Sean Wu, Wai-Mui Cheung, Ta-Yu Liu, Marvin Angelo E. Aberin, Ernesto Paas-Oliveros, Arian Escajeda, Edward Shih, Woan-Yuh Tarn, Yao-Ming Chang, Lan-Hsin Wang, Shu-Ping Wang

## Abstract

Histone H3K27 demethylase UTX (aka KDM6A) is mutated in many human cancers, suggesting its tumor suppressive role during cancer development. However, most tumors still express wild-type UTX/KDM6A and its function is not always linked to tumor suppression. Here, we present evidence of UTX/KDM6A’s role in sustaining tumor growth, revealing its function in tumor maintenance. We find that UTX/KDM6A sustains tumor cell cycling and survival via regulating DNA replication-associated transcriptional programs in a demethylase-independent manner. UTX/KDM6A can also interact with PARP1 and facilitate its recruitment to DNA lesions. Therefore, UTX/KDM6A depletion disrupts DNA replication and repair pathways, activating ATM–CHK2 and ATR–CHK1 signaling pathways and triggering S and G2/M checkpoints, leading to a pronounced defect in tumor growth. Analysis of human cancer xenograft models further demonstrates that knockdown of UTX/KDM6A by RNA-interference, rather than inhibition of its enzymatic activity via GSK-J4, shows potent anticancer effects. Dual inhibition of UTX/KDM6A and ATR further demonstrates synergistic anticancer activities. Our work highlights UTX/KDM6A as a potential therapeutic target for cancer treatment, especially when combined with ATR inhibition.

**Highlights:**

- UTX/KDM6A contributes to tumor maintenance by promoting the growth and survival of tumor cells
- Tumor cells rely on UTX/KDM6A to maintain DNA replication, cell cycling, and DNA damage repair
- UTX/KDM6A depletion triggers S and G2/M checkpoints via activating ATM–CHK2 and ATR–CHK1 signaling pathways
- Targeting UTX/KDM6A may prove to be an innovative strategy for cancer therapy, whether employed independently or in conjunction with ATR inhibitors.

**The Paper Explained:** *Problem:* The aggressive growth of tumors relies significantly on heightened proliferation rates and genomic instability, which necessitate robust DNA replication machinery and efficient DNA damage repair mechanisms for tumor cell survival and proliferation. UTX/KDM6A, a histone demethylase central to chromatin and epigenetic regulation, is commonly mutated in various human cancers. However, its role as a tumor suppressor or promoter remains unclear across different cancer contexts. This study delves into the potential tumor-maintaining role of UTX/KDM6A in cancer progression and tumorigenesis, establishing the mechanistic foundation for its tumor-promoting function.

*Results:* We uncover UTX/KDM6A’s crucial role in tumor maintenance via its participation in DNA replication and repair pathways. Surprisingly, we find that its histone demethylase activity is dispensable for these functions, implying an alternative role as a scaffold protein. Consequently, our findings suggest that targeting the entire UTX/KDM6A gene or protein, rather than inhibiting its enzymatic activity, holds promise as a therapeutic strategy for tumors dependent on its tumor-maintaining function.

*Impact:* This study unveils UTX/KDM6A’s multifaceted role in cancer progression, shedding light on its diverse contributions to tumorigenesis. Our findings suggest promising therapeutic strategies for cancer treatment, highlighting the importance of targeting UTX/KDM6A and its impact on DNA replication and repair pathways. These discoveries set the stage for further exploration of UTX/KDM6A-mediated mechanisms in clinical settings, indicating potential applications in future clinical trials and combination therapy strategies.

## Introduction

In eukaryotes, ensuring accurate genome duplication during cell division is paramount for maintaining genomic integrity (1). While the DNA replication machinery is inherently precise, it faces constant challenges from DNA damage induced by various intrinsic and extrinsic stresses (1, 2). Failure in this process can lead to the accumulation of DNA breaks and errors, culminating in genomic instability and driving cancer development. Despite being characterized by genomic instability, cancer cells rely heavily on robust DNA replication machinery (i.e. the replisome) and efficient DNA damage repair mechanisms for their growth and survival (1, 2).

Epigenetic alterations, including histone modifications and chromatin remodeling, play a central role in orchestrating DNA replication and responding to replication stress. Throughout S-phase progression, the spatiotemporal organization of DNA replication is intricately linked with the coordination of origin firing and higher-order chromatin structures (3). For instance, the EZH2-containing Polycomb Repressive Complex 2 (PRC2) binds to sites of DNA replication through the “parental” trimethylated H3K27 (H3K27me3) mark, thereby maintaining H3K27me3 at facultative heterochromatin and regulating genomic stability (4). Additionally, EZH2 mediates H3K27 methylation at stalled replication forks and contributes to the drug sensitivity of PARP inhibitor (PARPi) in BRCA2-deficient cancers (5). Beyond DNA replication, EZH2 plays a role in repair of DNA double-strand breaks (DSBs) (6) and transcriptional repression at enhancers and promoters (7–9). Therefore, the clinical relevance of EZH2 and the cognate H3K27me3 mark in tumorigenesis is established in various cancers (7, 8). Despite these insights, the roles of lysine-specific demethylases, particularly the KDM6 family (10, 11), in tumorigenesis, especially concerning DNA replication and repair, remain elusive.

UTX (gene name: KDM6A), a member of the KDM6 family, features six tetratricopeptide repeat (TPR) domains at the N-terminus and a catalytic Jumonji C (JmjC) domain at the C-terminus. This JmjC domain enables UTX/KDM6A to demethylate di- or trimethylated H3K27 (H3K27me2/me3), thereby counteracting the transcriptional repression exerted by polycomb proteins (10, 11). Additionally, UTX/KDM6A serves as a resident subunit of lysine methyltransferase MLL3/4 complexes (12). MLL3 and MLL4, also known as KMT2C and KMT2D respectively, deposit monomethylation at H3K4 (H3K4me1) in enhancer regions, delineating the enhancer chromatin landscape and enabling enhancer activation upon developmental cues (9, 12). Consequently, UTX/KDM6A is believed to coordinate H3K27 demethylation and H3K4 methylation on enhancers and/or promoters of target genes, thereby regulating cell and tissue differentiation through developmental gene activation. Importantly, recent evidence suggests that UTX/KDM6A can also modulate enhancer activation and gene expression through demethylase-independent mechanisms (13). These non-catalytic functions of UTX/KDM6A may stem from its direct interactions with chromatin-modifying enzymes, such as H3K4 mono-methyltransferases MLL3/4 (KMT2C/D), H3K27 acetyltransferases CBP/p300 (14, 15), and the chromatin-remodeling BAF (BRG1/BRM-associated factor) complex (16).

Numerous whole genome and exome sequencing studies have highlighted UTX/KDM6A as one of the most frequently mutated chromatin modifiers across various human cancers (17–19). These mutations, spanning a wide frequency range of 1% to 40%, are prevalent in diverse cancer lineages, such as bladder cancer, urothelial carcinoma, pancreatic cancer, esophageal squamous carcinoma, lung cancer, breast cancer, multiple myeloma, and T-cell acute lymphoblastic leukemia (T-ALL) (11, 17, 20). Consequently, UTX/KDM6A has long been regarded as a tumor suppressor during cancer development (11). However, intriguingly, aside from mutation variants (e.g., truncating mutations, missense mutations, in-frame insertions/deletions) that lead to UTX/KDM6A inactivation, gene amplifications of UTX/KDM6A have also been observed in numerous cancer types (19, 21). Furthermore, recent studies have demonstrated that pharmacological inhibition of UTX/KDM6A by a specific KDM6 inhibitor, GSK-J4, yields potent anticancer effects in neuroblastoma, ovarian cancer, and colorectal cancer (22–24). These findings strongly suggest a potential gain-of-function or tumor-maintaining role of UTX/KDM6A in tumorigenesis, particularly considering that a significant proportion of tumors consistently express wild-type UTX/KDM6A. However, it remains largely unclear whether UTX/KDM6A possesses the capacity to promote the growth and survival of tumor cells during cancer development and progression, and the underlying mechanisms warrant further investigation.

Here, we unveil an unprecedented mechanism wherein UTX/KDM6A serves a crucial role in supporting tumor cell growth and survival. Our study demonstrates that UTX/KDM6A regulates DNA replication-associated transcriptional programs and facilitates PARP1-dependent DNA repair through mechanisms independent of its demethylase activity. Depletion of UTX/KDM6A results in deficiencies in both DNA replication and repair pathways, thereby activating ATM– CHK2 and ATR–CHK1 signaling pathways, and inducing subsequent cell cycle arrest at the S and G2/M phases. These findings present compelling evidence for a novel role of UTX/KDM6A in tumor maintenance, underscoring its potential as an innovative target for cancer therapy.

## Results

### UTX/KDM6A is essential for tumor cell growth in multiple cancer types

Although UTX/KDM6A is reported to be mutated in multiple human cancers (11, 17, 20, 25) and is thought to function as a tumor suppressor during cancer development, wild-type UTX/KDM6A (wtUTX/KDM6A) still remains to be expressed in the majority of tumors and its function is not always linked to tumor suppression (21, 25, 26). We hypothesize that UTX/KDM6A is a highly pleiotropic factor in tumorigenesis and may play distinct roles in different cancer contexts or cancer stages. To test this hypothesis, we first analyzed and compared progression-free survival (PFS) in several cohorts of patients whose tumors express wild-type or mutant forms of *KDM6A* (Fig. S1). Despite the p-values not reaching significance, likely due to the limited number of patients with *KDM6A* mutations compared to those with wild-type *KDM6A*, we observed a negative correlation trend between the expression of wild-type *KDM6A* and PFS across various cancer types, such as non-small cell lung cancer (NSCLC), endometrial cancer, sarcoma and melanoma (Fig. S1B-D). These clinical observations suggest that UTX/KDM6A could be a risk factor in certain cancer conditions and its catalytic inhibition (or non-catalytic functions) may be advisable to be developed as a new cancer therapeutic strategy. To study whether UTX/KDM6A plays a role in supporting tumor growth, we analyzed the functional essentiality of UTX/KDM6A in tumor cell growth of multiple wtUTX/KDM6A-expressing cancer cell lines, including murine Lewis lung carcinoma (LL2) and colorectal carcinoma (CT26), human endometrial cancer (HeLa), NSCLC (NCI-H1975) and bone osteosarcoma (U2OS) cell lines. We conducted short hairpin RNA (shRNA)-mediated knockdown of UTX/KDM6A with two independent shRNA sequences (i.e., shUTX-7762 and shUTX-7763) in these cancer cell lines. These two independent shRNAs, which specifically target UTX/KDM6A for knocking down of gene expression, resulted in substantial reduction in UTX/KDM6A expression, as shown by Western blot analyses (Fig. 1A). Interestingly, colony formation assays showed that reduced UTX/KDM6A expression greatly attenuated the capability of tumor cell growth (Figs. 1B, 1C). Since the prolonged expression of UTX/KDM6A-shRNAs caused lethal phenotypes in tumor cells, we introduced a doxycycline (Dox)-inducible shUTX-7763 sequence for a stable and conditional knockdown of UTX/KDM6A in HeLa and NCI-H1975 cancer cell lines. We utilized NCI-H1975 and HeLa cell models due to the potential role of UTX/KDM6A in NSCLC and endometrial cancer development and progression (Figs. S1B, S1C). We performed a titration with Dox (50 to 250 ng/mL) induction and found that as little as 50 ng/mL was sufficient to induce UTX/KDM6A-targeting knockdown (Figs. 1D, 1E). Elevated levels of UTX/KDM6A-knockdown induced by higher dosages of Dox resulted in more pronounced growth inhibitory effects, confirming that selectively targeting UTX/KDM6A in tumor cells hinders both cell growth and survival (Fig. 1F). This supposition is further supported by the observation that Dox-induced UTX/KDM6A-knockdown dramatically suppressed HeLa and NCI-H1975 cancer cell proliferation in colony formation assays (Figs. 1G, 1H). Collectively, these results strongly underscore the indispensable functional role of UTX/KDM6A in supporting tumor cell proliferation across a spectrum of cancer lineages.

**Figure 1.**
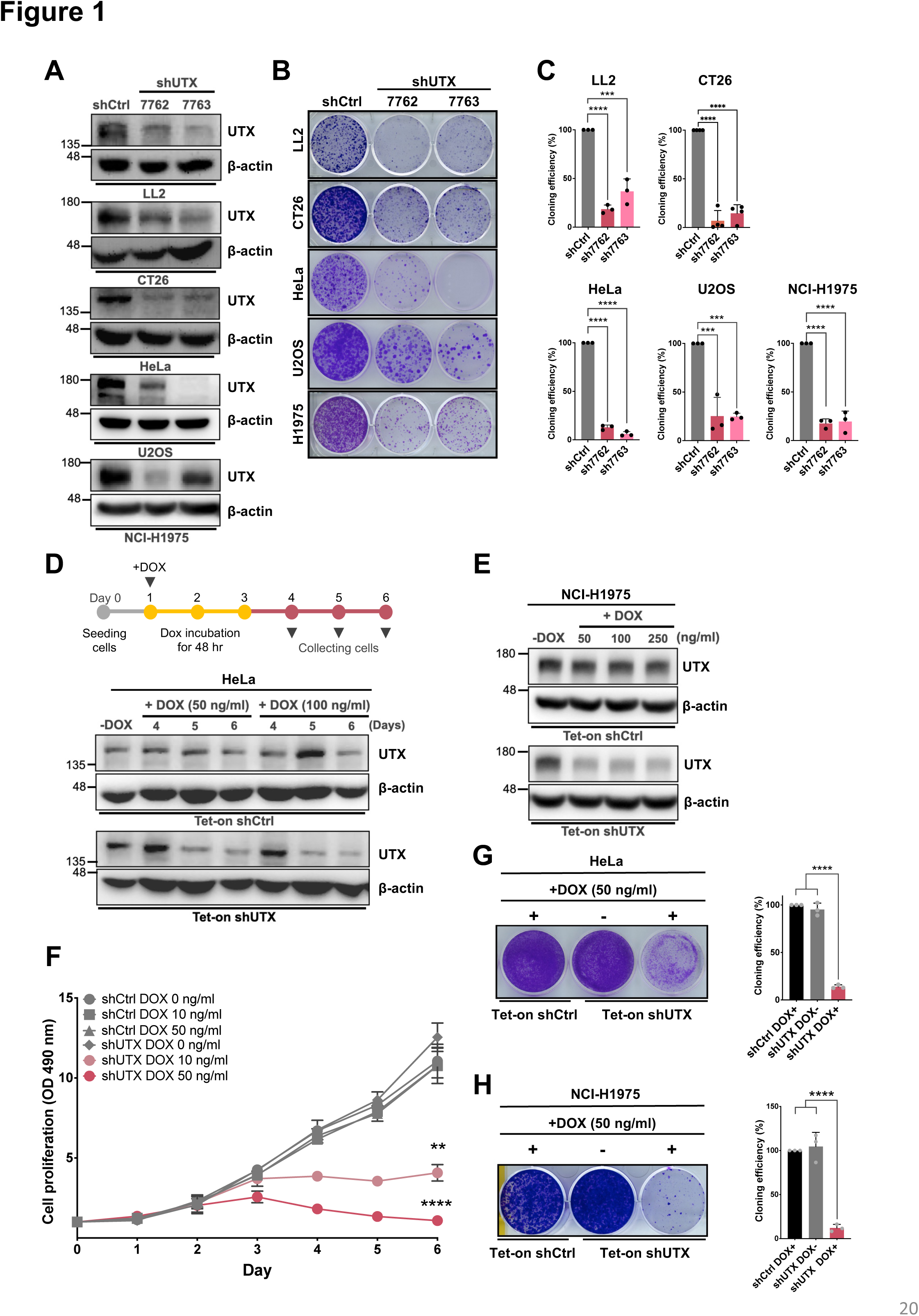
Unveiling a potential tumor-maintaining role of UTX in supporting tumor cell growth. (A) Western blot analysis was conducted with the specified antibodies on lysates from control (shCtrl) and UTX/KDM6A-shRNAs (shUTX-7762 or shUTX-7763)-transduced cancer cell lines. (B) Stable UTX/KDM6A-knockdown or control cells were subjected to colony formation assays, with representative plate images shown. (C) The colony formation ability of stable cell clones in (B) was assessed by crystal violet assays. Quantification is presented as mean `±` SEM for three independent experiments. (D) Top: A schematic illustrating the procedure for Dox-induced UTX/KDM6A-knockdown. Stably Dox-inducible (Tet-on) shUTX-expressing HeLa or control (shCtrl) cells were generated through lentiviral transduction. shRNA-mediated knockdown of UTX/KDM6A was induced by differential doses of Dox treatment as indicated. Lysates were collected at 4-, 5-, or 6-days post-Dox induction and subjected to Western blot analysis using the indicated antibodies. (E) Western blot analysis was performed using the specified antibodies on lysates obtained from Dox-induced UTX/KDM6A-knockdown NCI-H1975 cells. Lysates were collected at 3 days after Dox induction. (F) Cell viability was measured in Dox-induced UTX/KDM6A-knockdown HeLa or control cells treated with or without Dox. Cells were exposed to Dox treatment for 3 days and subjected to MTT assays as indicated time points. Data are presented as mean `±` SEM for three independent experiments. (G and H) Left: Colony formation assays were conducted for the indicated cells, HeLa for (G) and NCI-H1975 for (H), treated with or without Dox as the indicated doses. Representative images of the plates are shown. Right: The colony formation ability was assessed by crystal violet assays. Quantification is presented as mean `±` SEM for three independent experiments. **p < 0.01, ***p < 0.001, ****p < 0.0001 (Ordinary one-way ANOVA). Ctrl, control. Dox, doxycycline.

### UTX/KDM6A is essential for gene expression of the replisome that carries out DNA replication

To gain insight into the molecular mechanism by which UTX/KDM6A functionally supports tumor cell growth, we compared gene expression profiles in control and two independent UTX/KDM6A-knockdown HeLa cell clones using high-throughput RNA sequencing (RNA-seq). Both shRNAs-mediated UTX/KDM6A-knockdown resulted in comparable numbers of commonly downregulated and upregulated differentially-expressed genes (DEGs) (Figs. S2A, S2B). Since UTX/KDM6A is well-known to function as a transcription coactivator for gene regulation, we mainly focused on downregulated DEGs by UTX/KDM6A depletion in this study. Bioinformatics analyses of our RNA-seq data using KEGG pathway and gene set enrichment analysis (GSEA) showed that genes for DNA replication and repair pathways were downregulated by UTX/KDM6A depletion (Fig. 2A; Figs. S2C, S2D). Interestingly, we found that many of the DNA replication-associated factors, especially the components of replisome (2), may act directly downstream of UTX/KDM6A (Figs. 2B, 2C). The effects of UTX/KDM6A depletion on genes encoding the replisome components, including flap endonuclease 1 (FEN1), DNA ligase 1 (LIG1), minichromosome maintenance (MCM) subunits 3–7, replication factor C (RFC), proliferating cell nuclear antigen (PCNA), and replication protein A (RPA), were further confirmed in HeLa and NCI-1975 cell lines by RT-qPCR analyses (Figs. 2D, 2E; Figs. S2E, S2F).

**Figure 2.**
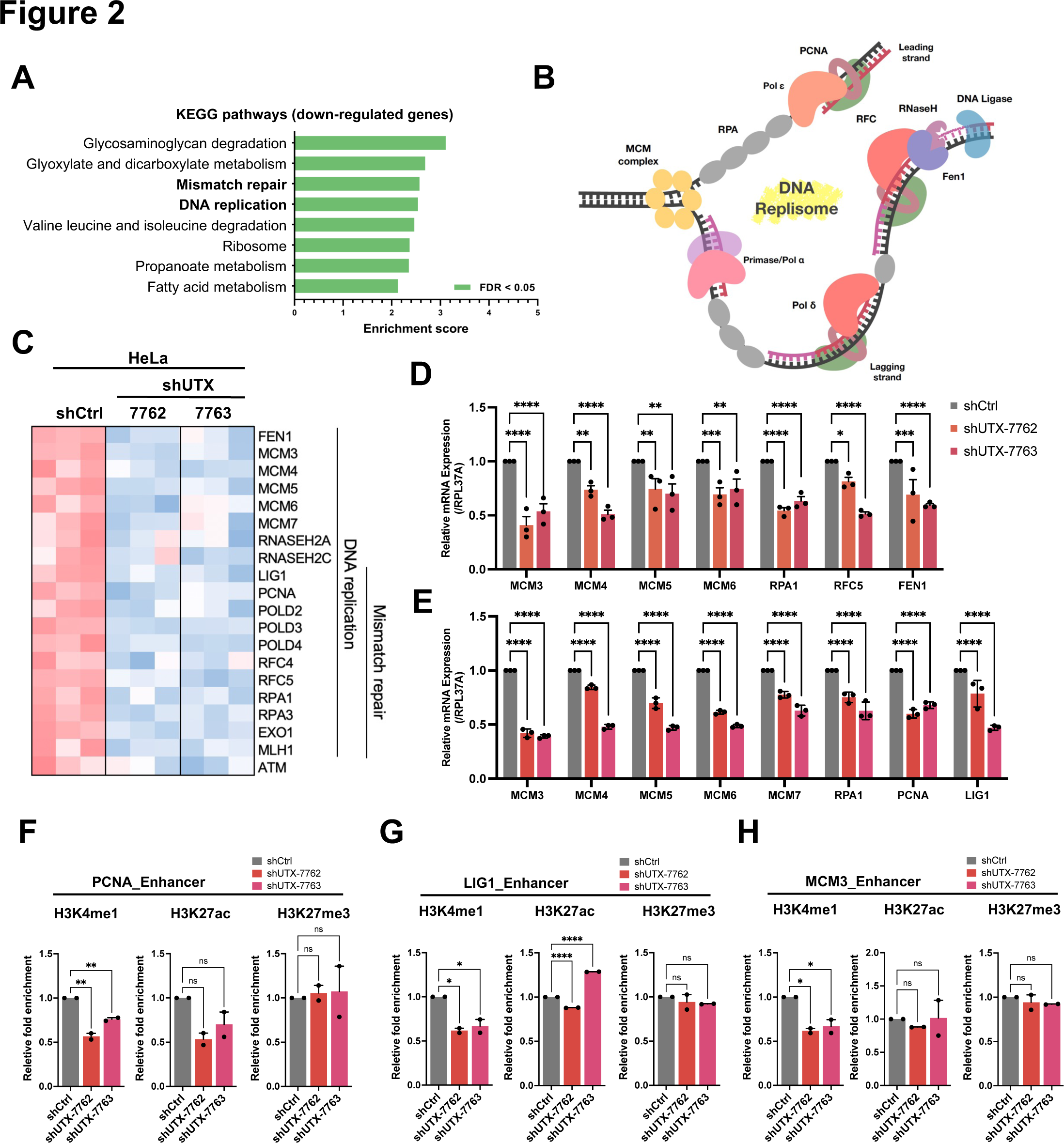
UTX/KDM6A orchestrates transcriptional programs associated with DNA replication and repair pathways. (A) Pathway enrichment analysis was conducted on genes (n = 3,349) downregulated by UTX/KDM6A depletion in HeLa cells, utilizing KEGG pathways. (B) A schematic representation delineates the components of the replisome converging on the replication fork. (C) Heatmaps illustrates genes associated with DNA replication and mismatch repair that exhibited downregulation in UTX/KDM6A shRNA-expressing HeLa cells compared to control cells. (D and E) RT-qPCR analysis was performed to assess mRNA levels of DNA replication-associated genes in HeLa (D) and NCI-H1975 (E) cells transduced with control (shCtrl) or UTX/KDM6A-shRNAs (shUTX-7762 or shUTX-7763). RPL37A served as an endogenous control. Data are represented as mean `±` SEM from three independent experiments. *p < 0.05, **p < 0.01, ***p < 0.001, ****p < 0.0001 (Two-way ANOVA). (F–H) Enrichments of enhancer histone marks (H3K4me1, H3K27ac, and H3K27me3) were analyzed at three representative gene enhancers in control (shCtrl) and UTX/KDM6A-shRNAs (shUTX-7762 or shUTX-7763)-transduced NCI-H1975 cells. The gene enhancers examined include *PNCA* (F), *LIG1* (G), and *MCM3* (H). Fold enrichments were normalized to H3, and relative fold enrichment was further normalized to the control cells (n = 2 biological replicates from 2 independent experiments). *p < 0.05, **p < 0.01, ****p < 0.0001, ns, no significant (Ordinary one-way ANOVA).

UTX/KDM6A is thought to exert its H3K27 demethylase activity to modulate the transition of chromatin signatures from the inactive/poised (i.e. H3K27me3) state to an active state marked by acetyl H3K27 (H3K27ac) (10, 11, 25). This involves UTX/KDM6A removing the methyl group from H3K27, providing a substrate for histone acetyltransferases p300 and CBP. However, UTX/KDM6A can also function as a scaffold, facilitating the recruitment and function of MLL4/KMT2D and p300 during the full activation of enhancer loci (13, 15). In this context, the H3K27 demethylase activity becomes dispensable for UTX/KDM6A-dependent gene activation. To further investigate whether UTX/KDM6A directly targets the enhancer and/or promoter regions of these putative target genes and activates transcription through a demethylase-dependent or independent mechanism, we first performed ChIP-qPCR analyses for H3K27me3, H3K27ac, H3K4me1 (enhancer) and H3K4me3 (promoter) marks. While UTX/KDM6A depletion significantly downregulated H3K4me1 at enhancer regions and H3K27ac at both enhancer and promoter regions to varying extents, it had no significant impact on H3K27me3 enrichment at either region (Figs. 2F-H; Figs. S2G-I). Notably, there was no significant alteration in H3K4me3 at target gene promoters following UTX depletion, despite a noteworthy decrease in H3K4me1 enrichment at these promoters (Figs. S2G-I). Furthermore, we utilized CUT&RUN to characterize H3K27me3 profiles in both control and UTX/KDM6A-knockdown NCI-H1975 cells. Interestingly, we observed no significant increase in H3K27me3 enrichment on the target gene loci (Figs. S2J-L). This further supports the notion that the demethylase activity of UTX/KDM6A is not essential for regulating genes associated with DNA replication. Additionally, treatment of NCI-H1975 cells with the GSK-J4 inhibitor significantly increased global H3K27me3 levels; however, no significant alteration in DNA replication-associated gene expression was observed (Figs. S2M, S2N). Hence, our data suggest that while UTX/KDM6A can regulate the activation of genes associated with DNA replication, the demethylase activity of UTX/KDM6A may not be essential for establishing and maintaining active enhancers and promoters of target genes. Instead, the activation of enhancer and promoter regions may necessitate the mono-methyltransferase activity of MLL3 or MLL4 (KMT2C/D), which could be associated with the non-catalytic function of UTX/KDM6A (15). Supporting this hypothesis, MLL4/KMT2D knockdown was found to reduce the expression of genes associated with DNA replication (Figs. S3A, S3B). Furthermore, we verified the interaction between UTX/KDM6A and MLL4/KMT2D in cells and identified a critical domain spanning residues 245–400, which encompass the C-terminal half of the TPR domains, that is essential for MLL4/KMT2D binding (Figs. S3C, S3D). To strengthen our conclusion that UTX/KDM6A collaborates with MLL4/KMT2D to promote the expression of genes associated with DNA replication, we utilized a publicly available dataset for NSCLC (27). We conducted multivariate least squares regression analyses to assess the correlation of gene expression with UTX/KDM6A, MLL4/KMT2D, and DNA replication-associated genes in NSCLC patient samples. Our analysis revealed a positive trend between the expression levels of DNA replication-associated genes and the expression levels of UTX/KDM6A and MLL4/KMT2D (Fig. S4). While the correlation is statistically significant in the 2D scatterplots (data not shown), it is worth noting that we may observe only a tendency due to the heterogeneity of the Cancer Genome Atlas samples in the 3D scatterplots.

### UTX/KDM6A depletion disturbs DNA replication, leading to a deceleration of S-phase and an arrest in G2/M-phase cell cycle progression

The eukaryotic replisome orchestrates all DNA replication processes, ensuring efficient and accurate chromosome replication (1, 2). However, replication can encounter obstacles, activating checkpoint surveillance systems that temporarily halt replication until issues are resolved (1, 2). Prompted by our data illustrating the impact of UTX/KDM6A depletion on DNA replication-associated transcriptional programs (Fig. 2), we investigated whether UTX/KDM6A deficiency in tumor cells triggers checkpoint activation, leading to cell cycle arrest. We conducted flow cytometry analysis of propidium iodide (PI) staining in Dox-induced UTX/KDM6A-knockdown HeLa and NCI-H1975 cells to examine the effect of UTX/KDM6A suppression on DNA synthesis and cell cycle progression. UTX/KDM6A-knockdown HeLa cells exhibited a robust G2/M arrest phenotype and underwent apoptosis (Figs. 3A, 3B), consistent with similar defects observed in UTX/KDM6A-knockdown NCI-H1975 cells under serum starvation (Figs. 3C, 3D). The occurrence of cell cycle arrest can also be confirmed by an increased level of phosphorylated histone H3 at serine 10 (H3S10P), a well-established marker of the mitotic index (28, 29). Therefore, we assessed alterations in cell cycling upon UTX/KDM6A loss by measuring the mitotic index in UTX/KDM6A-knockdown cells stained with an anti-H3S10P antibody. Consistently, a significant number of UTX/KDM6A-knockdown cells displayed abundant H3S10P foci compared to control cells (Figs. 3E, 3F), indicating impaired cell cycling due to UTX/KDM6A deficiency. To investigate the potential effects of UTX/KDM6A deficiency on S phase progression, we pulse-labeled Dox-induced UTX/KDM6A-knockdown and control HeLa cells with 5-bromodeoxoyuridin (BrdU) and analyzed S phase progression by FACS analysis. Control HeLa cells completed one S phase cycle (indicated by 4 N DNA content) and transitioned to the next cycle (indicated by 2 N DNA content) around 6-8 hours, whereas UTX/KDM6A-knockdown HeLa cells exhibited a pronounced S-phase delay, with a reduced population in early S-phase and an increased population in late S-phase compared to control cells (Figs. 3G, 3H). These findings suggest that UTX/KDM6A deficiency may induce S-phase delay and G2/M-phase arrest in tumor cells.

**Figure 3.**
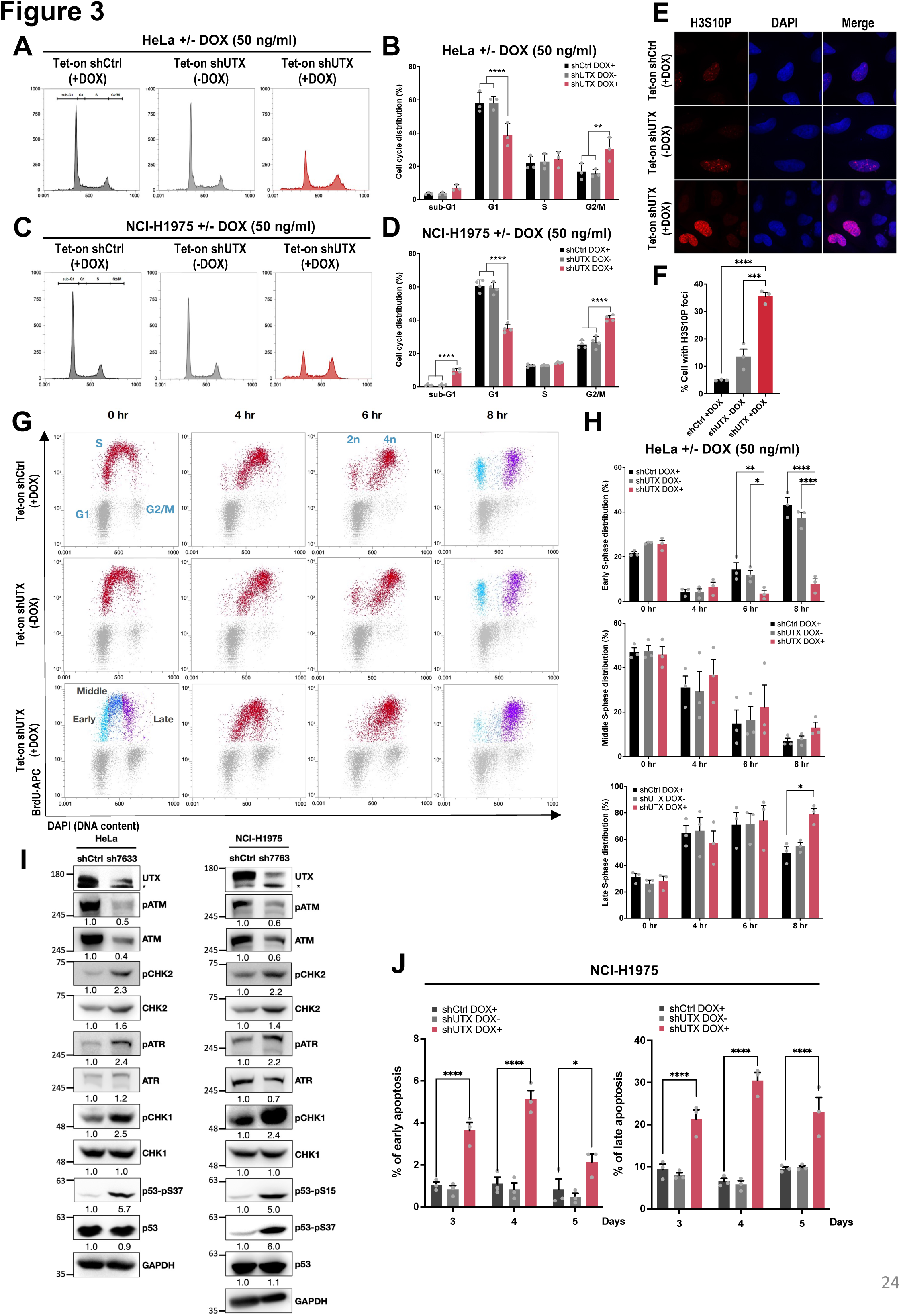
UTX/KDM6A deficiency disrupts S-phase and G2/M-phase cell cycle progression. (A) Flow cytometric analysis of the cell cycle was performed for Dox-inducible (Tet-on) shUTX-expressing HeLa or control (shCtrl) cells with or without Dox treatment. The analysis quantified sub-G1, G1, S, and G2/M phases from the PI-labeled histograms. Cells were treated with 50 ng/mL Dox for 72 h. (B) Quantification of cells in each phase is presented as mean `±` SEM for three independent experiments. (C) Flow cytometric analysis of the cell cycle was conducted for Dox-inducible (Tet-on) shUTX-expressing NCI-H1975 or control (shCtrl) cells with or without Dox treatment. Cell cycle analysis calculated sub-G1, G1, S, and G2/M phases from the PI-labeled histograms following serum starvation for 16 h. Cells were treated with 50 ng/ml Dox for 72 h. (D) Quantification of cells in each phase is shown as mean `±` SEM for three independent experiments. ****p < 0.0001 (Two-way ANOVA). (E) Immunostaining for histone H3 Ser10 phosphorylation (H3S10P) in Dox-inducible (Tet-on) shUTX-expressing HeLa or control (shCtrl) cells with or without Dox treatment (50 ng/mL). (F) Quantification of H3S10P signal intensity is presented as mean `±` SEM for three independent experiments. ***p < 0.001, ****p < 0.0001 (Two-way ANOVA). (G) Monitoring of defective S-phase progression in UTX/KDM6A-deficient cells. Dox-inducible (Tet-on) shUTX-expressing HeLa or control (shCtrl) cells were treated with 50 ng/mL Dox. Three days later, cells were pulse-labeled with BrdU and harvested at the indicated time intervals. Cells were stained with anti-BrdU monoclonal antibody and DAPI and analyzed by two-parameter FACS to measure DNA synthesis (BrdU-APC, y axis) and DNA content (DAPI, x axis). (H) Quantification of cells in early, middle, and late S phase is shown as mean `±` SEM for three independent experiments. (I) Western blot analysis was performed using the specified antibodies on lysates obtained from control (shCtrl) and UTX/KDM6A-shRNA (shUTX-7763)-transduced HeLa (left) or NCI-H1975 (right) cells. Numbers below the immunoblots panels indicate the densitometric values normalized to the respective GAPDH value. * denotes a non-specific band. (J) Cytotoxic effects of Dox-induced knockdown of UTX/KDM6A on NCI-H1975 cells were asseseed by flow cytometry with Annexin V-PI staining. Annexin V-positive and PI-negative populations represent cells in early apoptosis, while Annexin V and PI double-positive staining indicates cells in late apoptosis. *p < 0.05, **p < 0.01, ****p < 0.0001 (Two-way ANOVA). Ctrl, control. Dox, doxycycline.

### UTX/KDM6A depletion-induced replication stress triggers the activation of ATM–CHK2 and ATR–CHK1 signaling pathways

During DNA replication, DSBs can arise from the stalling, collapse or breakage of replication forks (30, 31). Consequently, DSB repair pathways are activated, either at replication forks or replicated chromatids, to ensure the completion of chromosome duplication (30, 31). The key DNA damage response (DDR) kinases ATM (ataxia-telangiectasia mutated) and ATR (ATM and rad3-related) play critical roles in checkpoint responses induced by DSB or replication stress, each with distinct DNA damage specificities (1, 30–32). ATM is primarily activated by DSBs, while ATR responds to a wide spectrum of DNA damage, encompassing various lesions that hinder DNA replication (1, 30–32). To study whether ATM and/or ATR-mediated checkpoint responses are evoked by UTX/KDM6A depletion and lead to S-phase delay and G2/M-phase arrest, we checked the phosphorylation status of ATM and ATR as well as the corresponding downstream effector kinases CHK2 and CHK1 in UTX/KDM6A-knockdown HeLa and NCI-H1975 cells. Interestingly, we observed a general decrease in overall protein levels of ATM in both UTX/KDM6A-knockdown HeLa and NCI-H1975 cells (Fig. 3I). These notable trends of downregulated ATM expression align with our RNA-seq results (Fig. 2C), suggesting that UTX/KDM6A may have the potential to regulate *ATM* gene expression as well. However, we could still detect a significant increase in CHK2 phosphorylation, implying that UTX/KDM6A loss may cause DSB accumulation and thereby trigger ATM–CHK2 signaling. Notably, we also detected a dramatic induction of ATR and CHK1 phosphorylation following UTX/KDM6A depletion (Fig. 3I). The induced phosphorylation of p53 at serine 15 (Ser15) and serine 37 (Ser37) provided additional confirmation of the activation of ATM and ATR in UTX/KDM6A-knockdown HeLa and NCI-H1975 cells (Fig. 3I). These data indicate that UTX/KDM6A loss may induce replication stress, leading to the subsequent accumulation of DSBs, which in turn triggers the activation of ATM and ATR and initiates subsequent DDR signaling. While UTX/KDM6A depletion led to a significant activation of the ATM–CHK2 and ATR–CHK1 checkpoint signaling pathways (Fig. 3I), our data revealed that UTX/KDM6A loss continued to elicit pronounced tumor cell apoptosis (Fig. 3J). Consequently, our data suggest that UTX/KDM6A depletion in tumor cells may not only impact DNA replication but also induce DNA breaks, thereby activating both ATR and ATM checkpoint signaling pathways.

### UTX/KDM6A plays a fundamental role in supporting DNA replication, thereby influencing cell cycling and survival in both tumor and non-tumorigenic/normal cells

Based on our observations, we propose that UTX/KDM6A likely plays a crucial role in supporting DNA replication, thereby influencing cell cycling and survival in both physiological and pathological contexts. Given the highly proliferative nature of tumor cells, it is conceivable that they may depend more on UTX/KDM6A compared to normal and non-tumorigenic cells. Consequently, we hypothesize that UTX/KDM6A depletion may result in fewer cytotoxic effects in normal cells and tissues. To test this hypothesis, we employed normal cell and animal models. Initially, we depleted UTX/KDM6A in human lung epithelial BEAS-2B cells and assessed its impact on DNA replication-associated transcriptional programs, cell cycling and the growth capacity of this non-tumorigenic cell line. Despite UTX/KDM6A depletion led to a downregulation of genes encoding the replisome components in BEAS-2B cells, the differences were modest and did not achieve statistical significance (Fig. S5A). Nonetheless, UTX/KDM6A depletion induced significant cell cycle arrest at S and G2/M phases in BEAS-2B cells, similar to that observed in NCI-H1975 cells (Fig. S5B). Notably, we observed a less prominent apoptotic population, as indicated by the sub-G1 cell fraction, in UTX/KDM6A-deficient BEAS-2B cells compared to UTX/KDM6A-deficient NCI-H1975 cells (Fig. S5B). This aligns with the reduced colony formation observed in UTX/KDM6A-deficient BEAS-2B cells, which was less evident compared to UTX/KDM6A-deficient NCI-H1975 cells (Figs. S5C, S5D). To explore the differential effects of UTX/KDM6A deficiency on tumor and non-tumorigenic cells, we examined ATM and ATR checkpoint pathways in control and UTX/KDM6A-deficient BEAS-2B cells. We consistently observed robust phosphorylation of CHK1 and CHK2 in UTX/KDM6A-deficient BEAS-2B cells (Fig. S5E), indicating a potential cell cycle arrest, similar to what was observed in Figure S5B. However, the induced phosphorylation of ATM and ATR, as well as the downstream effector p53 (Ser15 and Ser37), in UTX/KDM6A-deficient BEAS-2B cells were modest compared to that observed in UTX/KDM6A-deficient HeLa and NCI-H1975 cells (Fig. 3I; Fig. S5E). Hence, our results suggest that UTX/KDM6A depletion may induce a less pronounced DNA replication defect in non-tumorigenic BEAS-2B cells, as further supported by the modest activation of ATM and ATR signaling (Fig. S5E).

Our findings reveal that DNA replication defects observed in cells lacking UTX/KDM6A are consistent across transformed and non-transformed cells, albeit with differing degrees of severity. To gain further insights into the underlying mechanisms, we examined the expression levels of UTX/KDM6A in both transformed NCI-H1975 and non-transformed BEAS-2B cells. Remarkably, the expression of UTX/KDM6A in NCI-H1975 cells significantly exceeds that in BEAS-2B cells (Fig. S5F), supporting the notion that tumor cells may rely on higher levels of UTX/KDM6A protein to sustain their highly proliferative state. We further explored the significance of UTX/KDM6A and its demethylase function in supporting tumor cell growth. For this purpose, we employed a Dox-inducible UTX/KDM6A overexpression system. Colony formation assays revealed that both wild-type and a demethylase-dead mutant (HE/AA) of UTX/KDM6A effectively rescued the reduced colony formation observed in UTX/KDM6A-deficient BEAS-2B cells to a similar extent (Figs. S5G, S5H). Although the rescued phenotypes were less apparent in UTX/KDM6A-deficient NCI-H1975 cells, potentially due to incomplete compensation by UTX/KDM6A overexpression for the deficiency caused by UTX/KDM6A-shRNAs, we found that overexpression of either wild-type or the demethylase-dead mutant of UTX/KDM6A significantly promoted colony formation in UTX/KDM6A-deficient NCI-H1975 cells compared to the no overexpression control (Figs. S5I, S5J). These findings underscore the non-essential role of UTX/KDM6A demethylase activity in tumor cell growth through DNA replication regulation, revealing a similar regulatory mechanism in both transformed and non-transformed cells with varying levels of UTX/KDM6A dependency.

To further explore the physiological role of UTX/KDM6A in DNA replication *in vivo*, we focused on the evolutionarily conserved endoreplication process, wherein cells undergo repeated DNA replication cycles without cell division (33, 34). A prime model for studying this phenomenon is the giant salivary glands of *Drosophila melanogaster* where cells undergo DNA replication (S) and gap (G) phases without mitosis, resulting in a large nucleus with approximately 1350 copies of DNA content (35). A defective endoreplication process in *Drosophila* salivary glands leads to reduced nuclear size (33, 34, 36, 37). Therefore, the nuclear size of *Drosophila* salivary glands serves as a reliable indicator of DNA replication status. In the homozygous null mutant of *Drosophila* UTX/KDM6A homolog (dUtx^-/-^), we observed that the nuclear size of salivary glands was smaller than that of wild-type *Drosophila* (Figs. S6A, S6B). These findings indicate impaired endoreplication in the absence of dUtx, although the effects of dUtx loss on the nuclear size of salivary glands were less dramatic compared to the loss of other cell cycle regulators (37, 38). To further assess the effects of dUtx loss on normal cell growth and survival, we examined the essentiality of dUtx in *Drosophila* wing development. In *Drosophila melanogaster*, adult wings develop from monolayered epithelia in the pouch region of larval wing imaginal discs (39). In *Drosophila* larvae carrying homozygous null mutations of dUtx, we observed a slight increase in the activation of the apoptosis effector caspase Dcp-1 in the wing pouch epithelia compared to the wild-type control (Figs. S6C, S6D). These data suggest that although loss of dUtx may affect DNA replication process in normal *Drosophila* cells, dUtx deficiency only causes minor cellular apoptosis (∼1.4% of the wing epithelial tissue), consistent with our observations in human BEAS-2B cell line.

Taken together, our data suggest that loss of UTX/KDM6A has relatively subtle effects in normal cells and tissues. However, the perturbations associated with its depletion in tumor cells provide insights into its potential role as a drug target for cancer therapeutics (see details below).

### UTX/KDM6A depletion also impairs HR and NHEJ-mediated DNA damage repair

Our findings reveal that UTX/KDM6A depletion not only significantly activated the ATM– CHK2 and ATR–CHK1 checkpoint signaling pathways but also continued to induce pronounced tumor cell apoptosis (Figs. 3I, 3J). This prompted us to explore whether UTX/KDM6A may also play a role in the repair of DNA replication stress-induced DSBs. This may reflect the substantial cellular apoptotic effects induced by the loss of UTX/KDM6A. For this purpose, we employed hydroxyurea (HU), a DNA replication stress agent known for depleting cells of deoxyribonucleotide triphosphates (dNTPs), thereby inducing DNA replication stress (40). We exposed both control and UTX/KDM6A-knockdown HeLa cells to 2mM HU treatment, followed by immunofluorescence staining of histone H2AX phosphorylation at serine 139 (γH2AX). This histone modification serves as the primary cellular biomarker in response to DSB damage (32). As expected, depletion of UTX/KDM6A led to a significant increase in γH2AX foci, further accentuating their appearance following HU-induced DNA replication stress (Figs. 4A, 4B). The induction of γH2AX foci was more pronounced when we induced DSBs in HeLa cells by exposure to DNA-damaging agents, such as ionizing radiation (IR) (Figs. 4C, 4D). These data suggest that UTX/KDM6A may also contribute to DSB repair, whereas loss of UTX/KDM6A promotes severe DNA damage events. To further investigate whether UTX/KDM6A contributes to DSB repair pathways, we induced DSBs in HeLa cells by a low dose of γ-radiation (2 Gy) and measured the clearance of γH2AX foci as a readout of DSB repair. Although IR induced similar amount of γH2AX foci in control and UTX/KDM6A-knockdown cells at early time points after the treatment, these foci disappeared more slowly in UTX/KDM6A-knockdown cells than in the control cells at late time points post-IR exposure (Figs. 4C, 4D). Taken together, these results indicate that UTX/KDM6A is essential for DDR, including focus formation, checkpoint activation, and the overall DSB repair.

**Figure 4.**
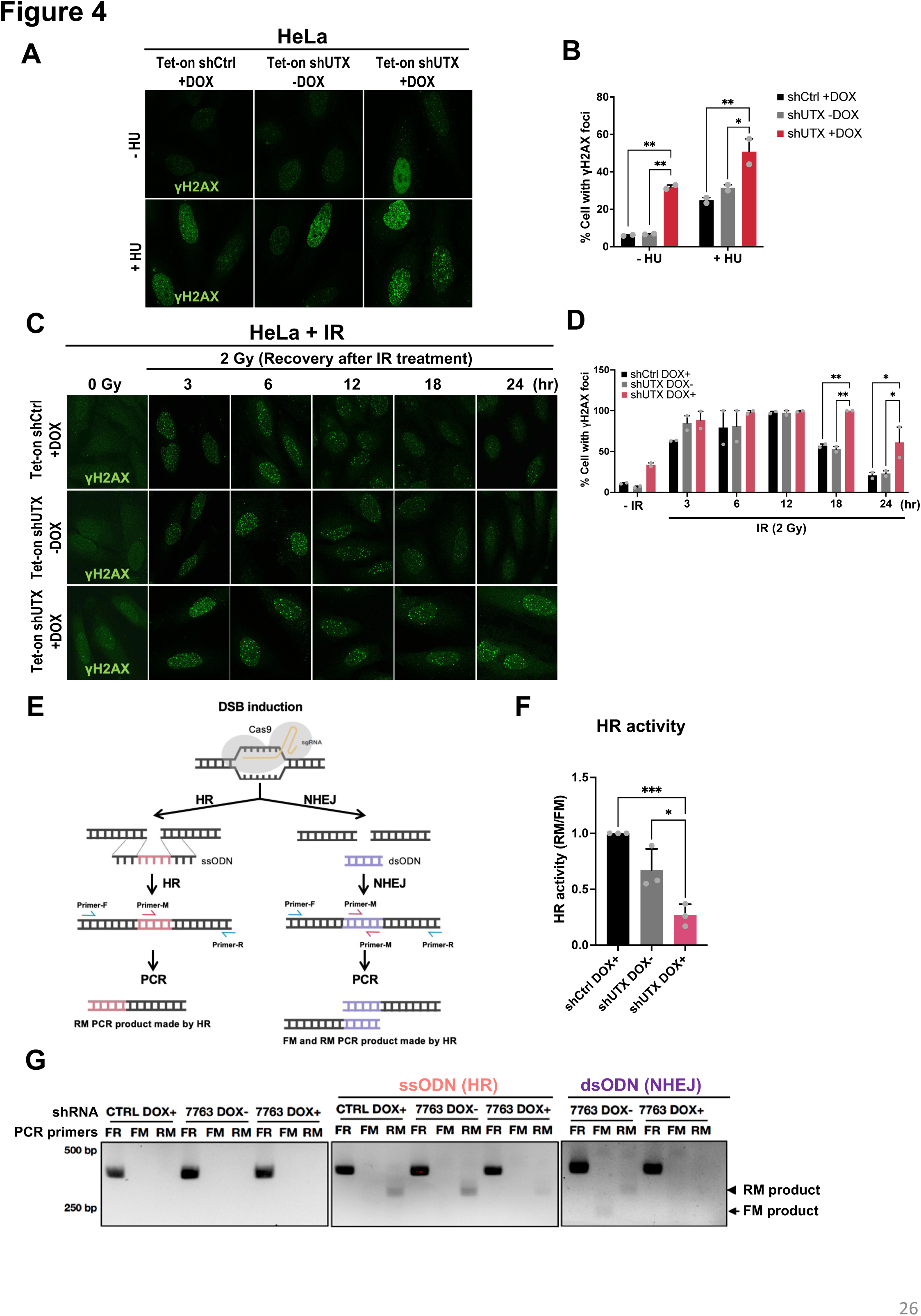
UTX/KDM6A deficiency impairs DDR and DSB repair. (A) Dox-inducible (Tet-on) shUTX-expressing HeLa cells treated with or without Dox (50 ng/mL) and 0 or 2mM of hydroxyurea (HU) for 72 h were stained for γH2AX. (B) Quantification of γH2AX signal intensity is presented as mean `±` SEM for three independent experiments. (C) Dox-inducible (Tet-on) shUTX-expressing HeLa or control (shCtrl) cells treated with or without Dox (50 ng/mL) and 0 or 2 Gy γ-radiation, followed by recovery for indicated time points, were stained for γH2AX. (D) Quantification of γH2AX signal intensity is presented as mean `±` SEM for three independent experiments. *p < 0.05, **p < 0.01 (Two-way ANOVA). (E) The schematic diagram illustrates a unique DSB-repair monitoring system. Site-specific DSBs were introduced via CRISPR/Cas9-mediated genomic DNA cutting, followed by integration of the unique marker sequence (incorporated in ssODN or dsODN) via HR or NHEJ pathways. PCR analysis was then employed to quantitatively evaluate the marker sequence in the genomic DNA. ODN, oligodeoxynucleotide. (F) Genomic DNA was collected 24 h after CRISPR/Cas9-mediated site-specific DSBs. The activity of HR or NHEJ was determined by PCR analysis. PCR products amplified with different primer sets (i.e., FR, FM, RM) were analyzed by electrophoresis in a 2% agarose gel. The arrow indicates the FM PCR product, and the arrowhead indicates the RM PCR product. (G) HR activity was quantified by qPCR analysis. Quantification of HR activity is presented as mean `±` SEM for three independent experiments. *p < 0.05, ***p < 0.001, (Ordinary one-way ANOVA). Ctrl, control. Dox, doxycycline.

The repair of DSBs in mammalian cells relies on two primary mechanisms: homologous recombination (HR) and non-homologous end joining (NHEJ) (32). To investigate and quantify the effects of UTX/KDM6A deficiency on DSB repair pathways, we adapted a method based on CRISPR/Cas9-induced oligodeoxynucleotide (ODN)-mediated HR and NHEJ (41). This approach involves incorporating ODN at the CRISPR/Cas9-induced DSB sites, where single-stranded ODN (ssODN) serves as a donor template for HR pathway, while blunt-ended double-stranded ODN (dsODN) can be integrated into DSB sites by NHEJ (Fig. 4E). By designing specific primer sets (i.e., FR, FM, RM), semi-quantitative or real-time polymerase chain reaction (PCR) can be used to detect the activity of HR or NHEJ. We utilized this method to evaluate the cellular activity of HR and NHEJ in both control and UTX-knockdown HeLa cells. Interestingly, we found that UTX/KDM6A deficiency greatly reduced the efficiency of both HR and NHEJ repair mechanisms in cells (Figs. 4F, 4G). These findings suggest that UTX/KDM6A has the potential to regulate DSB repair by modulating the cellular activity of HR and/or NHEJ. Moreover, our data imply that, besides its role in DNA replication-associated gene regulation, UTX/KDM6A may also contribute to the repair of DNA replication stress-induced DSBs.

### UTX/KDM6A interacts with PARP-1 and facilitates its recruitment to DNA damage sites

To dissect the molecular mechanism through which UTX/KDM6A regulates DSB repair, we directed our focus towards poly(ADP-ribose) polymerase 1 (PARP-1) for several compelling reasons: (i) PARP-1 serves as a well-recognized DNA damage sensor, pivotal in DNA base excision repair (BER) and repair of DNA single-strand breaks (SSBs) and DSBs (32); (ii) PARP-1 activation occurs in response to DNA replication stress, operating at normal and stalled replication forks (32); (iii) the interplay between PARP-1 and PRC2, which counteracts UTX/KDM6A’s function in H3K27me3 modification, has implications in DSB repair (6, 42, 43). Consequently, we hypothesized a potential interplay between UTX/KDM6A and PARP-1 in DDR and DSB repair. To evaluate this hypothesis, we established U2OS cell lines expressing GFP-tagged PARP-1 (GFP-PARP-1) and investigated whether UTX/KDM6A influences the recruitment of GFP-PARP-1 to DNA lesions (Fig. S7A). Utilizing laser micro-irradiation to induce DNA damage, we monitored the real-time recruitment of GFP-PARP-1 to DNA damage sites (Fig. S7B). Our observations revealed rapid recruitment of GFP-PARP-1 to laser-induced DNA damage, peaking at approximately 1-minute post-irradiation (Figs. 5A-C). Notably, GFP-PARP-1 recruitment was significantly impaired upon shRNA-mediated UTX/KDM6A knockdown (Figs. 5A-C; Fig. S7C), suggesting a dependency of PARP-1 recruitment on UTX/KDM6A or its H3K27 demethylase activity. Supporting this notion, *in vitro* binding assays with purified proteins demonstrated a direct interaction between UTX/KDM6A and PARP-1 (Fig. 5D). Furthermore, through the generation of deletion mutants, we identified both the N-terminal TPR and C-terminal JmjC domains on UTX/KDM6A as crucial for its interaction with PARP-1 (Figs. 5E, 5F), indicating the necessity of multiple UTX/KDM6A domains for PARP-1 interaction. The physical association between PARP-1 and UTX/KDM6A was further confirmed using co-immunoprecipitation assays (Fig. 5G). Interestingly, we could also detect a complex containing PARP-1, UTX/KDM6A, and MLL4/KMT2D (Fig. 5H), suggesting that the UTX/MLL4 (KDM6A/KMT2D) complex may contribute to PARP-1 recruitment to DNA damage sites and the subsequent chromatin relaxation (44) (see detailed discussion below). While we could not exclude the possibility that the H3K27 demethylase activity may facilitate PARP-1 recruitment to DNA lesions, our findings underscore the indispensable role of UTX/KDM6A in PARP1-dependent DDR and DSB repair.

**Figure 5.**
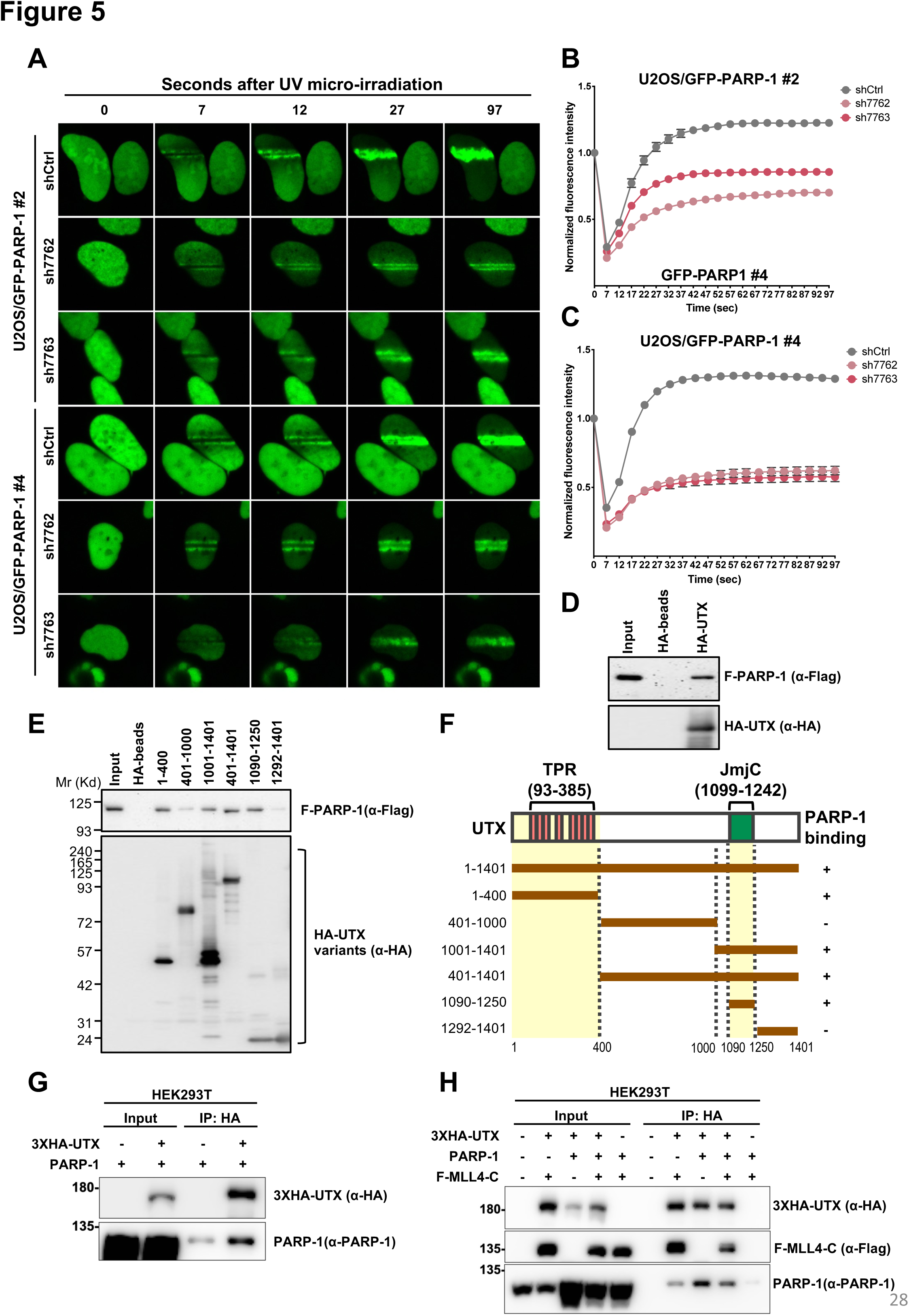
UTX/KDM6A facilitates PARP-1 recruitment to DNA damages sites during DNA damage response. (A) Two independent U2OS subclones expressing GFP-PARP-1 (U2OS/GFP-PARP-1 #2 and #4) with stable transduction of shCtrl, shUTX-7762, or shUTX-7763 were subjected to laser micro-irradiation and monitored using time-lapse microscopy. Representative images at 0, 7, 12, 27, and 97 seconds are shown after laser micro-irradiation. (B and C) The accumulation of GFP-PARP1 on the laser damage tracks in U2OS/GFP-PARP-1 #2 (B, n = 22) or #4 (C, n = 25) cells was quantified and plotted. Increased fluorescence on the damage tracks is plotted over time. (D and E) Direct binding of purified Flag-tagged PARP-1 (F-PARP-1) to full-length (D) or deletion fragments (E) of HA-tagged UTX (HA-UTX) immobilized on anti-HA beads (monitored by antibodies as indicated). (F) Schematic representation and summary of the interaction between PARP-1 and various deletion fragments of UTX/KDM6A. Ctrl, control. (G) PARP-1 interacts with UTX/KDM6A in vivo. HEK293T cells were co-transfected with PARP-1 and 3XHA–UTX (or an empty vector) for 36 h. Subsequently, cell lysates were immunoprecipitated (IP) with anti-HA or mouse IgG and subjected to immunoblotting using the specified antibodies. (H) Interactions among PARP-1, UTX/KDM6A, and MLL4/KMT2D in cells. HEK293T cells were co-transfected with PARP-1, 3XHA–UTX, Flag-MLL4-C (C-terminal region, amino acids 4507-5537), or the corresponding empty vectors for 36 h. Following transfection, cell lysates were immunoprecipitated (IP) with anti-HA or mouse IgG and subjected to Western blot analysis using the specified antibodies.

### UTX/KDM6A inactivation through gene silencing, rather than inhibition by GSK-J4, demonstrates potent anticancer efficacy in human cancer xenograft mouse models

To investigate the potential of UTX/KDM6A blockade in suppressing tumor growth *in vivo*, we employed human cancer xenograft mouse models. Dox-inducible shUTX-expressing NCI-H1975 or control (shCtrl) cells were subcutaneously implanted into the backs of BALB/c nude female mice. Upon entry into the exponential growth phase, mice received a diet and water supplemented with Dox throughout the experiment. We observed that diminished UTX/KDM6A expression via UTX/KDM6A-shRNAs effectively inhibited NCI-H1975 tumor growth, with significant reduction in tumor size and weight (Figs. 6A-C). To further evaluate whether UTX/KDM6A catalytic inhibition could phenocopy the response of NCI-H1975 cells to the whole UTX/KDM6A depletion, we treated mice bearing the control (shCtrl) NCI-H1975 cells with the GSK-J4 inhibitor. In contrast to the whole UTX/KDM6A depletion, GSK-J4 exhibited no significant effect on the growth of NCI-H1975 tumors (Fig. S8A). We demonstrated that both shUTX-mediated gene silencing and GSK-J4-mediated UTX/KDM6A catalytic inhibition led to a global increase in H3K27me3 levels in NCI-H1975 cells (Figs. S8B, S8C), suggesting that the tumor-maintaining function of UTX/KDM6A may be uncoupled from its H3K27 demethylase activity. Hence, our findings suggest that the loss of UTX/KDM6A’s tumor-maintaining function, potentially independent of its H3K27 demethylase activity, disturbs DNA replication and repair processes in tumor cells, thereby impeding tumor cell proliferation and inducing tumor cell apoptosis.

**Figure 6.**
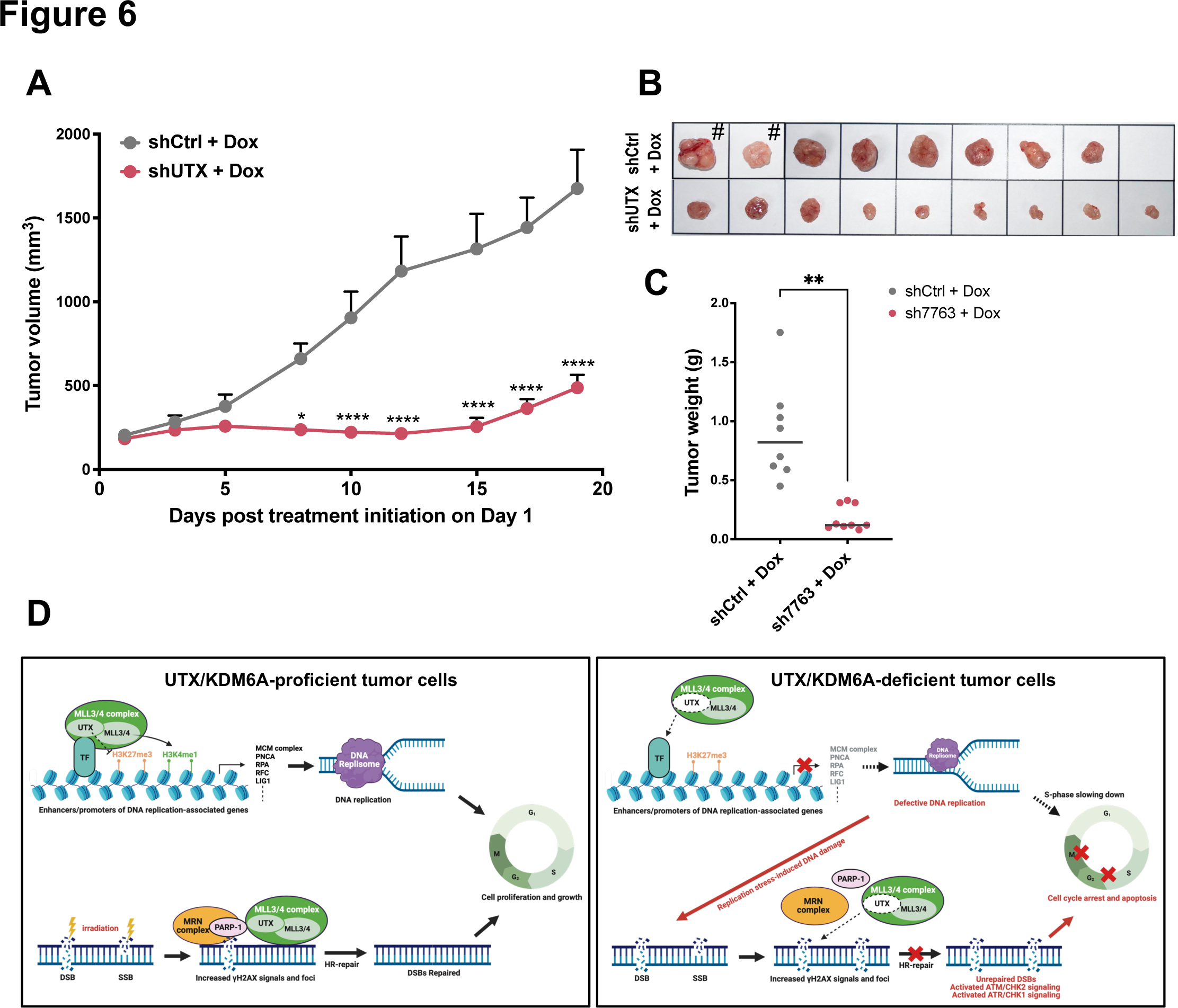
The tumor-maintaining function of UTX/KDM6A supports tumor growth *in vivo*. (A) Tumor growth rates of Dox-inducible (Tet-on) shUTX-expressing NCI-H1975 or control (shCtrl) cell lines upon Dox treatment as indicated in xenograft assays (shCtrl, n=8; shUTX, n=9). Data are represented as means `±` SEM. *p < 0.05, ****p < 0.0001 (Two-way ANOVA followed by Bonferroni correction). (B) Gross pathological photographs at necropsy of the NCI-H1975 tumors that arose in mice treated with Dox-induced shCtrl or shUTX (shCtrl, n=8; shUTX, n=9). #, Two mice in the shCtrl group were euthanized prior to the scheduled endpoint (Day 17 post-treatment) due to their tumor sizes surpassing 2000 mm³. (C) The weight of each tumor in the indicated tumor groups. **p < 0.01 (Unpaired t test with Welch’s correction). Ctrl, control. Dox, doxycycline. (D) Proposed model illustrating the tumor-maintaining functions of UTX/KDM6A in the regulation of DNA replication and repair pathways to support tumor cell growth. In UTX/KDM6A-proficient tumor cells, UTX/KDM6A, independently of its H3K27 demethylase activity, plays a key role in activating DNA replication-associated gene expression, probably by physically bridging the MLL3/4 (KMT2C/D) complex to target gene enhancers and promoters. UTX/KDM6A may also facilitate the interaction between PARP-1 and the MLL3/4 (KMT2C/D) complex (and probably the MRN complex) to DNA damage sites in response to intrinsic and extrinsic genomic stresses. In UTX/KDM6A-deficient tumor cells, loss of UTX/KDM6A results in defective DNA replisome function, which may slow down the S-phase progression, induce replication stress, and lead to DNA damage accumulation. Due to the critical roles of UTX/KDM6A in DDR and DNA repair (e.g., HR-repair), UTX/KDM6A-deficient tumor cells fail to repair the replication stress-induced DNA damage. Consequently, loss of UTX/KDM6A results in the accumulation of substantial DNA damage, triggering DNA replication and DSB checkpoint signaling pathways, provoking cellular apoptosis, and ultimately leading to severe tumor cell death. Created with BioRender.com.

### Targeting the ATR pathway renders tumor cells more vulnerable to UTX/KDM6A blockade

While our findings demonstrate that loss of UTX/KDM6A provokes tumor cell apoptosis, we postulate that the activated ATR–CHK1 signaling may still mitigate the detrimental effects of replication stress, thereby protecting tumor cells from apoptosis. This is supported by our colony formation data, which indicate a small fraction of tumor cells survive from UTX/KDM6A depletion (Figs. 1A-C). Preclinical studies have highlighted that tumor cells with compromised checkpoint or DNA repair mechanisms may exhibit heightened sensitivity to ATR inhibition (45, 46). Consequently, several ATR inhibitors have entered clinical trials, holding promise for cancer therapeutics (45, 46). Building upon these observations, we hypothesized that targeting the ATR pathway could sensitize tumor cells to UTX/KDM6A depletion. Indeed, our data reveal that while both the UTX/KDM6A-shRNA and the ATR inhibitor AZD6738 significantly impair colony formation in NCI-H1975 cells, the combined treatment of UTX/KDM6A-shRNA and AZD6738 exerts a greater tumor cell-killing effect (Figs. S9A-C). Notably, the combined treatment resulted in not only fewer colonies but also smaller average colony size in UTX/KDM6A-deficient NCI-H1975 cells treated with AZD6738 (Figs. S9A, S9C), suggesting that dual inhibition of UTX/KDM6A and ATR synergistically inhibits tumor cell growth and survival. Additionally, robust synergistic induction of apoptosis was observed in NCI-H1975 and HeLa cells when shRNA-induced UTX/KDM6A knockdown was combined with AZD6738, as indicated by CellEvent Caspase-3/7 assays (Figs. S9D-F). In line with our cell line-based studies, which demonstrate strong synergistic anti-tumor effects with dual inhibition of UTX/KDM6A and ATR, the efficacy of this combination therapy for cancer therapeutics has been evaluated in NCI-H1975 xenograft mouse models (Fig. S9G).

## Discussion

### UTX/KDM6A: Orchestrating Tumor Maintenance through DNA Replication, Cell Cycling, and Genome Stability

Our study elucidates a mechanistic rationale for considering UTX/KDM6A as a key molecular target to assess therapeutic efficacy in tumors dependent on its tumor-maintaining function. We observe its pivotal role in orchestrating DNA replication-associated transcriptional programs and regulating the PARP1-dependent DNA repair pathway. Given the heightened proliferative demand and genomic instability characteristic of tumor cells, these distinctive functions of UTX/KDM6A likely play a crucial role in supporting tumor cell growth and survival. Consistent with this notion, we demonstrate that UTX/KDM6A knockdown results in impaired DNA replication and repair pathways, culminating S and G2/M cell cycle arrest. Despite the substantial activation in the ATM–CHK2 and ATR–CHK1 checkpoint signaling pathways following UTX/KDM6A knockdown, cellular apoptosis is significantly induced, leading to extensive tumor cell death (Fig. 6D). Building upon the therapeutic potential inferred from these observations, we evaluate the effects of shRNA-mediated UTX/KDM6A inactivation in human NSCLC xenograft models. Our results underscore the potent anticancer efficacy of selectively targeting UTX/KDM6A, particularly against tumors reliant on its tumor-maintaining function.

### H3K27 Demethylase-independent Functions of UTX/KDM6A in DNA Replication and Repair Pathways

Intriguingly, our investigation into the functions of UTX/KDM6A has unveiled a fascinating aspect: its role in facilitating DNA replication and repair pathways appears to be independent of its H3K27 demethylase activity. Specifically, our ChIP-qPCR analysis has shown a significant decrease in H3K4me1 levels at the enhancers and promoters of genes associated DNA replication upon UTX/KDM6A depletion, while the levels of H3K27me3 at these gene loci remained largely unaffected. Further investigation has illuminated that the N-terminal motif, encompassing the TPR domain, is indispensable for UTX/KDM6A’s interaction with MLL4/KMT2D. This interaction is pivotal for orchestrating the recruitment of MLL4/KMT2D to enhancer sites (15, 47), thereby establishing active enhancer landscapes crucial for the activation of target genes associated with DNA replication. Therefore, it is highly possible that the N-terminal motif of UTX/KDM6A is specifically responsible for mediating this demethylase-independent function, facilitating the direct recruitment of MLL4/KMT2D to enhancer and promoter regions for the transcription of genes crucial for DNA replication. However, a comprehensive understanding of the binding motifs governing the reciprocal interaction between UTX/KDM6A and MLL4/KMT2D, as well as their roles in DNA replication-associated transcriptional programs, necessitates further exploration and detailed characterization.

Second, our findings point out another demethylase-independent function of UTX/KDM6A in controlling PARP1-mediated DNA repair pathways. We demonstrated that UTX/KDM6A interacts directly with PARP-1 through multiple regions, including the TPR and JmjC domains, potentially facilitating PARP-1 recruitment to DNA damage sites. This is consistent with recent research showing that MLL3/4 (KMT2C/D) regulates DSB repair by directly binding to DNA lesions, where it mediates H3K4 methylation, chromatin relaxation, and recruitment of secondary DDR factors, such as NBS1 and MDC1 (44). The recruitment of MLL3/4 (KMT2C/D) to DNA lesions is facilitated by Argonaute 2 (AGO2) and small noncoding DNA damage response RNAs (DDRNAs), which are processed by RNA polymerase II (Pol II), Drosha, and Dicer (44, 48–50). Initially, DSBs trigger the primary recruitment of PARP-1 and the MRE11–RAD50–NBS1 (MRN) complex, potentially coordinating the recruitment of Pol II, Drosha and/or Dicer to nearby regions (48, 49, 51). Based on these findings, it is conceivable that UTX/KDM6A could play a role in mediating the interaction among MLL3/4 (KMT2C/D), PARP-1, the MRN complex, and AGO2 to initiate and expand DDR signals (Fig. 6D). The co-immunoprecipitation data (Fig. 5H), demonstrating the presence of a cellular complex containing UTX/KDM6A, MLL4/KMT2D, and PARP-1, aligns with this proposed model. Since the JmjC catalytic domain of UTX/KDM6A contributes to the UTX/KDM6A-PARP1 interaction, further investigation is needed to understand whether and how the H3K27 demethylase activity of UTX/KDM6A regulates chromatin relaxation during DNA repair. While the N-terminal region of UTX/KDM6A appears essential for the interaction with MLL3/4 (KMT2C/D) and PARP-1, the specific motifs involved in these interactions requires additional verification.

### UTX/KDM6A Loss Leads to Downregulation of Replisome Components, Resulting in DNA Replication Slowdown and Replication Stress

Accurate DNA replication requires various factors, including nucleotide pools (dNTPs), replisome components, histones, and histone chaperones. Limitation in these essential factors often result in replication progression slowdown and eventual replication stress (30, 52). Our studies pinpoint a crucial role of UTX/KDM6A in transcriptionally activating genes encoding key replisome components, such as MCM subunits, LIG1 and PCNA. We propose that the downregulation of these replisome components due to UTX/KDM6A loss may impede DNA replication speed, thereby promoting replication stress (Fig. 6D). Consistent with this notion, our data reveal that loss of UTX/KDM6A induces a notable increase in endogenous DNA damage events and triggers hyperphosphorylation of ATR and CHK1 checkpoint kinases (Fig. 3I). During DNA replication, repair of replication stress-induced DNA damage relies on HR utilizing the sister chromatid DNA as a template (1, 2, 30, 31). Consequently, the observed activation of the DNA replication checkpoint likely serves to stabilize stalled replication forks, halt cell cycle progression, and facilitate HR-dependent DNA damage repair for replication fork restart (1, 2, 30, 31). Interestingly, our findings shed light on a potential mechanism underlying the role of UTX/KDM6A in influencing HR repair (Fig. 4F). We propose that while UTX/KDM6A loss-induced replication stress triggers the DNA replication checkpoint, defective HR repair machinery may fail to completely solve replication defect. As a result, loss of UTX/KDM6A leads to DNA damage accumulation, culminating in cellular apoptosis and profound tumor cell death (Fig. 6D).

### The Tumor Maintaining Function of UTX/KDM6A: Implications for Tumor Growth across a Spectrum of Cancers

Detrimental mutations in *KDM6A*, encoding UTX, are prevalent in various human cancers, notably urothelial bladder carcinoma, pancreatic cancer, and certain blood cancers. Corresponding genetically engineered mouse models (GEMMs) support a tumor suppressive role of UTX/KDM6A in these cancers (11, 17, 19, 25). Despite the highest mutation frequency being observed in urothelial bladder carcinoma, with rates reaching up to 40%, *KDM6A* mutations are found in fewer than 10% of the most common human cancers (e.g., breast, lung, prostate, and colorectal cancer) and other cancer types (Fig. S1A). These clinical observations suggest that UTX/KDM6A may play a tumor-maintaining and promoting role in human cancers independent of UTX/KDM6A mutations, potentially contributing to tumor growth and malignancy (Figs. S1B-E). Indeed, emerging research highlights UTX/KDM6A’s involvement in cancer development and progression across various contexts and stages (21, 25, 26). For instance, UTX/KDM6A supports the oncogenic function of estrogen receptor α (ERα) in breast cancer development (53, 54) and is crucial for breast cancer metastases via regulating mesenchymal-epithelial transition (MET) (55). Notably, the tumor-maintaining and promoting role of UTX/KDM6A in breast cancer malignancy is dependent on its H3K27 demethylase function, as evidenced by the suppressive effect of GSK-J4 on breast cancer stem cells (BCSCs) and MET-associated transcriptional programs (55, 56). GSK-J4 has also shown promise in glioma treatment, particularly in diffuse intrinsic pontine gliomas (DIPGs), where somatic H3K27 mutations lead to global H3K27 hypomethylation supporting glioma development (26, 57). Furthermore, GSK-J4 acts as a radiosensitizer, enhancing the efficacy of radiotherapy in DIPGs by reversing H3K27 hypomethylation states on DNA damage repair genes and augmenting radiation-induced DNA damage (58). Thus, the efficacy of GSK-J4 in tumor treatment is context-dependent, influenced by H3K27 hypomethylation states and specific cancer contexts.

Importantly, our findings suggest that the H3K27 demethylase activity of UTX/KDM6A is largely dispensable for its tumor-maintaining and promoting function. While this activity may contribute to chromatin relaxation and/or PARP1-dependent damage response and repair, our data emphasize the non-catalytic functions of UTX/KDM6A in DNA replication-associated gene activation and the recruitment of PARP-1 to DNA lesions. Notably, UTX/KDM6A mRNA levels positively correlate with those of DNA replication-associated genes in various cancers, including sarcoma, melanoma, endometrial and lung cancer (Table S1). Elevated levels of these DNA replication-associated gene signatures strongly correlate with tumor development (Fig. S10). Consistent with these findings, our human cancer xenograft models demonstrate that whole UTX/KDM6A gene depletion, rather than inhibition via GSK-J4, exerts a more potent antitumor effect (Fig. S8A). Moreover, simultaneous inhibition of UTX/KDM6A and ATR demonstrates synergistic anticancer activities (Fig. S9G). Collectively, our data suggest that targeting the entire UTX/KDM6A gene or protein may serve as a new cancer therapeutic strategy for human tumors requiring its tumor-maintaining and promoting function. This approach should be tailored to specific patient cohorts, and the co-expression of *UTX*/*KDM6A* and DNA replication-associated genes could serve as a crucial component of future clinical trials.

## Methods

### Cell Culture

The human NCI-H1975 non-small cell lung cancer (NSCLC), HeLa endometrial cancer, U2OS bone osteosarcoma, HEK293T embryonic kidney, and mouse LL2 Lewis lung carcinoma, CT26 colorectal cancer cell lines were cultured in specific media: RPMI-1640 for NCI-H1975 and LL2, DMEM for HeLa, HEK293T and CT26, and McCoy’s 5A for U2OS. These media were supplemented with 10% FBS (Gibco) and 1% penicillin/streptomycin (Gibco), and cells were maintained at 5% CO_2_ and 37°C in a humidified incubator.

### Generation of UTX/KDM6A-knockdown Cell Lines

To achieve stable knockdown of UTX/KDM6A in cells, two shRNAs targeting KDM6A (shUTX-7762 and shUTX-7763) were cloned into the shRNA lentiviral knockdown vector pLKO.1 pure, a gift from Bob Weinberg (Addgene plasmid # 8453; http://https://www.addgene.org/8453; RRID: Addgene_8453). For doxycycline-induced knockdown of UTX/KDM6A, the Tet-on shUTX (shUTX-7763) oligonucleotides, designed as described in a previous report (59), were cloned into the EZ-Tet-pLKO-Puro vector, a gift from Cindy Miranti (Addgene plasmid # 85966; https://www.addgene.org/85966; RRID: Addgene_85966). shRNA targeting sequences (with RNAi consortium ID) are as follows: shUTX-7762 (TRCN0000107762), shUTX-7763 (TRCN0000107762), and shMLL4 (TRCN0000013138).

### Clonogenic Survival Assay

To assess cell survival and proliferation, cells were seeded at a density of 5 x 10^3^ to 1 x 10^4^ cells per well in 6-well plates, with triplicates for each condition. For the doxycycline inducible-shUTX expressing cells or the control (shCtrl) cells, cells were cultured in the presence or absence of doxycycline (Sigma-Aldrich, Cat#D9891). The culture media were refreshed after 3 days, and colonies became visible after 10 to 14 days of incubation. Colonies were then stained with 0.25% crystal violet (Sigma-Aldrich, Cat#C0775) and washed with PBS before air-drying. The stained colonies were dissolved by adding lysing solution (100 mM sodium citrate [pH4.2], 50% ethanol) and incubated for 30 min at room temperature. Optical density (OD) was measured at 590 nm using the SpectraMax 190 Microplate reader (Molecular Devices). The cloning efficiency was determined by normalizing the absorbance of the treated samples to that of the untreated controls.

### Western Blot Analysis

Cells were lysed in ice-cold RIPA buffer (50 mM Tris-HCl [pH7.5], 5 mM EDTA, 0.1% SDS, 0.5% sodium deoxycholate, 1% NP-40, 150 mM NaCl) supplemented with both proteinase inhibitors (Roche, Cat#4693132001) and phosphatase inhibitors (Roche, Cat#4906837001) for 15 min on ice. Following lysis, cell lysates were sonicated for 4 cycles (30s on/30s off, high setting) using a Bioruptor (Diagenode), followed by a 15-min incubation on ice. The supernatant containing the lysate was collected by centrifugation at 15,000 rpm for 20 min and quantified using Pierce™ BCA Protein Assay Kits (Thermo Fisher Sc., Cat#23227). Equal amounts of protein were mixed with SDS sample buffer and boiled at 95°C for 5 min. Subsequently, proteins were separated by SDS-PAGE, transferred onto polyvinylidene fluoride (PVDF) membranes, and detected by immunoblotting using the specified antibodies. Immunoblots were imaged using the UVP BioSpectrum Auto Imaging System (Analytik Jena). The list of antibodies and their sources can be found in supplemental table S2.

### Cell Viability Assay

The inducible-shUTX expressing cells or the control (shCtrl) cells were seeded at a density of 1 x 10^3^ cells per well in a 96-well plate, with triplicates for each condition. After 18 h, cells were cultured in the presence or absence of doxycycline. At specified time points, cells were treated by directly adding MTS reagents (BioVision, Cat#K300) and incubated for 60 min at 37°C. Cell proliferation was assessed by measuring absorbance at 490 nm and calculated as fold change by normalizing to the absorbance measured of the first day.

### Quantitative Reverse Transcription PCR (RT-qPCR)

Total RNA was extracted using the RNAspin Mini Kit, and cDNA was synthesized from 1 μg RNA using PrimeScriptTM RT Master Mix kit (Takara, Cat#RR036A) following the manufacturer’s protocols. Quantitative real-time PCR (qPCR) was performed using the Power SYBR Green PCR Master Mix (Applied Biosystems, Cat#4368708) and detected on the QuantStudio 5 Real-Time PCR system (Applied Biosystems). The housekeeping gene *RPL37A* served as an internal control for normalization. Target gene expression levels were normalized to *RPL37A* mRNA expression levels and calculated using the deltaCt (ΔCt) method. The experiments were performed in biological triplicate, and the data were presented as mean ± SEM. The primers utilized in RT-qPCR can be found in supplemental table S3.

### Chromatin Immunoprecipitation (ChIP) Assay

ChIP assays were performed as previously described with minor modifications (60). Briefly, cells were fixed with 1% (vol/vol) formaldehyde for 10 min at room temperature and following by quenching with 125 mM glycine. Fixed cell pellets were collected by centrifugation at 3,000 rpm for 5 min and washed twice with ice-cold PBS. Subsequent steps were carried out at 4°C. To extract chromatin, cell pellets were lysed by Buffer 1 (50 mM HEPES, 1 mM EDTA [pH 8.0], 0.5% NP-40, 0.25% Triton X-100, 150 mM NaCl) supplemented with 2x complete EDTA-free Protease Inhibitor Cocktails (PIC, Roche, Cat#04693132001), and incubated for 10 min with rotation. After incubation, cells were pelleted by centrifugation at 15,000 rpm for 10 min, resuspended in Buffer 2 (10 mM Tris-HCl [pH 8.0], 5 mM EDTA, 2.5 mM EGTA, 200 mM NaCl, and 2x PIC freshly added), and rotated for 10 min. Following centrifugation, cell pellets were resuspended in Buffer 3 (10 mM Tris-HCl [pH 8.0], 5 mM EDTA, 2.5 mM EGTA, and 2x PIC freshly added), and rotated for an additional 10 min. To achieve a final concentration of 0.5% N-lauroylsarcosine, 10% N-lauroylsarcosine was added and incubated for 10 min with rotation. For sonication, lysates were transferred to the Covaris microtube (15x19 mm) and sonicated for 20 min using a program with 140W intensity, 200 burst time, and 5% duty cycle in the S220 Focused Ultrasonicator (Covaris). Under this condition, chromatin was fragmented to sizes ranging from 150 to 500 bp. After sonication, insoluble material was removed by centrifugation at 15,000 rpm for 20 min, and the sheared chromatin was collected. The sheared chromatin was then incubated with 2 µg of specified antibodies in IP buffer (15 mM Tris-HCl [pH 6.8], 1% Triton X-100, 150 mM NaCl, and 1x PIC) overnight, followed by recovery through binding to Dynabeads protein G (Thermo Fisher Sc., 10004D) for an additional 3 h. After sequential washing steps with RIPA I buffer (50 mM HEPES, 10 mM EDTA [pH 8.0], 0.5% sodium deoxycholate, 0.5% NP-40, 500 mM LiCl) and RIPA II buffer (50 mM HEPES, 10 mM EDTA [pH 8.0], 0.5% sodium deoxycholate, 0.5% NP-40, 50 mM LiCl), each repeated three times for 5 min with rotation, beads were resuspended in TES buffer (50 mM Tris-HCl [pH 8.0], 10 mM EDTA, 1% SDS) and incubated at 65°C for 15 min for elution. Beads were removed using a DynaMag-2 Magnet (Thermo Fisher Sc., 12321D). The eluate was then incubated with 0.2 mg/mL RNase A at 37°C for 20 min and subsequently with 0.2 mg/mL Proteinase K at 55°C overnight. After reverse crosslinking, DNA was purified using QIAquick PCR Purification Kit (Qiagen). The experiments were performed in biological duplicate and the data were presented as mean ± SEM. The primers utilized in ChIP-qPCR can be found in supplemental table S3.

### Flow Cytometry Analysis

The inducible-shUTX expressing cells or the control (shCtrl) cells were harvested via trypsinization and subsequently washed with ice-cold PBS. Following centrifugation, cells were fixed with 70% ice-cold ethanol by gently vortexing, followed by overnight incubation at -20°C. The cell pellets were then collected by centrifugation and resuspended in ice-cold PBS containing 10 µg/mL propidium iodide (PI), followed by a 30-min dark incubation at 37°C. The stained cells were analyzed using the Attune CytPix Flow Cytometer (Thermo Fisher Sc.).

### Bromodeoxyuridine (BrdU) Incorporation Assay

The inducible-shUTX expressing cells or the control (shCtrl) cells were incubated with doxycycline (50 ng/mL) for 72 h, followed by treatment with 10 µM 5-bromodeoxoyuridin (BrdU) for 30 min. After BrdU incorporation, cells were replenished with culture media for recovery and collected at specified time points using eBioscienceTM BrdU Staining Kit for Flow Cytometry APC (Invitrogen, Cat#8817660042), following the manufacture’s protocols. The BrdU incorporated cells were then analyzed using Attune CytPix Flow Cytometer.

### Immunofluorescence Assay

The inducible-shUTX expressing cells or the control (shCtrl) cells were plated on glass slides and treated with doxycycline (50 ng/mL) or hydroxyurea (2mM) for 72 h. To induce DNA damage, cells were exposed to ionizing irradiation (IR) as indicated doses and then fixed by 4% paraformaldehyde at room temperature for 10 min. Subsequently, cells were permeabilized twice by incubating in 0.2% Triton X-100/PBS buffer at room temperature for 10 min, followed by blocking in buffer (0.5% BSA, 0.2% Triton X-100, PBS) at room temperature for 30 min. Cells were then incubated with the indicated primary antibodies overnight at 4°C, washed three times with 0.2% Triton X-100/PBS buffer (5 min each), and incubated with fluorescence-conjugated secondary antibodies at room temperature for 3 h. The nuclei of fixed cells were stained with DAPI and ProLong™ Diamond Antifade Mountant (Thermo Fisher Sc., Cat#P36961) to minimize photobleaching and achieve optimal signal brightness. Images were captured using the Zeiss LSM780 microscope (Carl Zeiss), and the intensity of γH2AX-positive cells was quantified using the ImageJ software. Total cell numbers were obtained from at least 150 cells per experiment. The experiments were performed in biological triplicate, and the data were presented as mean ± SEM.

### Quantitative Assessment of HR and NHEJ Activities

The quantitative assessment of HR and NHEJ activities was performed as previously described (41). Briefly, cells expressing the inducible shUTX or the control (shCtrl) were seeded at a density of 5 x 10^4^ per well in triplicate a in 6-well plate. After 48 h of doxycycline (50 ng/mL) treatment, cells were transfected with the CRISPR-Cas9 plasmids (1 µg), together with the ssODN (20 pmol) or dsODN (25 pmol), using Lipofectamine 2000 reagents (Invitrogen, Cat#11668019) in accordance with the manufacturer’s instructions. At 24 h post-transfection, cells were harvested for genomic DNA extraction using the PureLink™ Genomic DNA Mini Kit (Invitrogen, Cat#K1820-02). For HR activity assessment, qPCR was performed using 100 ng of genomic DNA, 200 nM primer sets (i.e., FR, FM, RM), and Power SYBR^TM^ Green PCR Master Mix (Invitrogen) in a final volume of 20 µl. The qPCR assays followed these thermal cycling conditions: 1 cycle at 95 °C for 2 min; 35 cycles at 95°C for 30s, 53°C for 30s, 72°C for 40s; and 1 cycle at 72 °C for 10 min. The cycle threshold (Ct) value of FR served as the internal control for normalization with the RM product. The experiments were performed in biological triplicate, and the data were presented as mean ± SEM.

### Laser Micro-irradiation

GFP-PARP-1 stable cell lines, with or without doxycycline-induced UTX knockdown, were seeded in Nunc™ Lab-Tek™ II Chambered Coverglass (Thermo Fisher Sc., Cat#155409). After treatment with 20 µM Hoechst 33342 for 10 min, cells were placed on the heated stage (37°C) of a laser-scanning confocal microscope (LSM780). DNA damage was induced along a 0.2–1 μm-wide region across the nucleus by exciting the Hoechst 33342 dye using a 20 mW, 405 nm laser line. The laser output was adjusted to 100% power with 200 iterations to create localized DNA damage, utilizing a Plan-Apochromat 100x/1.4 oil DIC immersion objective. GFP fluorescence imaging was captured every 5 sec post-excitation. GFP fluorescence intensity was quantified in the laser exposure region and normalized against the initial image (pre-excitation). The experiments were performed in biological triplicate, and mean along with standard error were plotted based on a minimum of 15 cells.

### Protein interaction assays

For co-immunoprecipitation assays, HEK293T cells were transiently transfected with plasmids encoding PARP-1, Flag-tagged MLL4-C (amino acids 4507-5537), 3XHA-tagged full-length of UTX, and/or UTX deletion variants as indicated. Post-transfection, cells were harvested, and the cell pellets were lysed using lysis buffer (50 mM Tris-HCl [pH 7.4], 150 mM NaCl, 1 mM EDTA, 0.1% Triton X-100, 0.5 mM DTT, 1 mM PMSF), supplemented with 1 x PIC, for 30 min on ice. The lysates were subsequently clarified by centrifugation at 15,000 rpm for 30 min, and the resulting supernatant was collected. This supernatant was then incubated with magnetic beads conjugated with IgG for 1 h at 4°C to perform pre-clearing. Following pre-clearing, the lysate was incubated with anti-HA magnetic beads (MBL International) for 4 h at 4°C to facilitate the immunoprecipitation of the protein of interest. Subsequently, the anti-HA magnetic beads were subjected to three washes with lysis buffer, each lasting 5 min, at 4°C to remove non-specifically bound proteins. Finally, the bound proteins were eluted from the magnetic beads by adding sample buffer. For in vitro binding assays, recombinant proteins were generated using the Bac-to-Bac Baculovirus Expression Systems (Thermo Fisher Sc.). These recombinant proteins were then immobilized on anti-HA magnetic beads and subsequently incubated, as indicated, with purified factors in BC150 buffer (20 mM Tris-HCl [pH 7.5], 0.2 mM EDTA, 150 mM KCl, 20% glycerol) supplemented with 0.1% NP40, 3% BSA and 1 mM PMSF. Reactions were conducted at 4°C for 4 h. Following thorough washing with binding buffer, the bound proteins were resolved by SDS-PAGE and subjected to immunoblotting using the specified antibodies for analysis.

### In Vivo Mouse Xenograft Study

The NCI-H1975/shCtrl and NCI-H1975/shUTX cells were seeded into coated T-175 flasks with 50 mL of culturing medium and incubated in an atmosphere of 5% CO_2_ at 37°C. The cells underwent two passages per week for expansion and were tested to ensure they were free of mycoplasma before being harvested for animal implantation. Female BALB/c nude mice, 6 weeks of age, received subcutaneous implants of viable NCI-H1975/shCtrl and NCI-H1975/shUTX tumor cells, each at a concentration of 2 x 10^6^ cells with 50% Matrigel per mouse in a 0.1 mL volume of culture medium, into the right flank. Tumor volumes were measured using calipers to determine length and width, with length representing the largest tumor diameter and width representing the perpendicular tumor diameter. Tumor volumes was calculated using the prolate ellipsoid formula: tumor volume = length x (width)^2^ x 0.5. Eight days post tumor implantation, animals with grouped mean tumor volumes between 121 and 124 mm^3^ began treatment, designated as Day 1. On Day 1, animals were provided with doxycycline-containing chow (Envigo Bioproducts Inc., Cat#TD.01306) to induce UTX/KDM6A knockdown. Concurrently, doxycycline hyclate (Sigma-Aldrich, Cat#D9891) was formulated in water for injection at 2.5 mg/mL and orally (PO) administered to animals twice weekly at 25 mg/kg with a dosing volume of 10 mL/kg. Doxycycline treatment commenced on Day 1 and continued untill study completion. GSK-J4 (MedChemExpress, Cat#HY-15648B) was formulated in 5% DMSO with 5% Solutol in PBS at 5 mg/mL and intraperitoneally (IP) administered at 50 mg/kg at a dosing volume of 10 mL/kg daily for 8 consecutive days (QD x 8 days). AZD6738 (also known as Ceralasertib, MedChemExpress, Cat#HY-19323) was formulated in 10% 2-hydroxyl-propyl-b-cyclodextrine in PBS solution and PO administered at 50 mg/kg at 10 mL/kg dosing volume daily daily for 8 consecutive days (QD x 8 days). Animals were monitored daily for humane endpoints, and those found in a moribund state were humanely sacrificed. Data on body weight (data not shown) and tumor volume of control and treatment groups over time were plotted using GraphPad Prism. Significant reduction in tumor volume was determined by the indicated p-value for statistical significance comparing treatment groups to the vehicle control using two-way ANOVA followed by Bonferroni correction. Mice were sacrificed 3–4::Jweeks after tumor implantation, or when tumor size exceeded 2000::Jmm^3^. Tumor weight at the conclusion of experiments were plotted by comparing treatment groups with the vehicle control using GraphPad Prism software and analyzed using Unpaired t test with Welch’s correction. Error values are presented as SEM values. All procedures involving animals, including housing, experimentation, and disposa, adhered to the Guide for the Care and Use of Laboratory Animals: Eighth Edition (National Academy Press, Washington, D. C., 2011). Pharmacology Discovery Services Taiwan, Ltd. (PDS), have been fully accredited by the Association for Assessment and Accreditation of Laboratory Animal Care International (AAALAC International) in 2014, 2017 and 2020 (#001553). Animal care and use protocols were reviewed and approved by the IACUC at PDS.

### Apoptosis Analysis

The NCI-H1975/shCtrl and NCI-H1975/shUTX cells were harvested at indicated time points and stained with PI and AnnexinV-FITC (Strong Biotech Corp., Cat#AVF250) according to the manufacture’s protocols. The stained cells were analyzed by Attune CytPix Flow Cytometer. The experiments were conducted in biological triplicate, and the data were presented as mean ± SEM. For assessing the synergistic effects of ATR inhibitor AZD6738 on UTX/KDM6A loss-induced cell apoptosis, apoptosis was evaluated in NCI-H1975 or HeLa cells using CellEvent Caspase-3/7 Green ReadyProbe Reagent (Invitrogen, Cat#R37111) as per the manufacturer’s instructions. Briefly, the inducible-shUTX expressing or the control (shCtrl) cells were cultured with and without doxycycline and/or AZD6738 on glass slides. Cells were then incubated with CellEvent Caspase-3/7 Green ReadyProbes Reagent for 30 min at 37°C and fixed in 4% paraformaldehyde for 10 min at room temperature. After permeabilization twice with 0.2% Triton X-100/PBS buffer for 10 min at room temperature, cells were incubated in blocking buffer (0.5% BSA, 0.2% Triton X-100, PBS) for 30 min at room temperature. The nuclei of fixed cells were stained by DAPI. Images were captured using the Zeiss LSM780 microscope, and the intensity of apoptotic cells (i.e., green fluorescence-positive cells) was quantified using ImageJ measurements. Mean and standard error for a minimum of 150 cells were counted.

### Statistical Methods and Data Resources

Statistical significance of experiments in this paper were assessed using GraphPad Prism with appropriate methods as indicated. The raw reads of RNA-seq analysis of transcriptional changes in the UTX-knockdown (shUTX-7762 and shUTX-7763) versus the control (shCtrl) HeLa cells in this paper have been deposited to the NCBI SRA database with the accession ID of PRJNA1031333.

## Acknowledgements

We thank Yu-Ling Lee, Kun-Yuan Lin, Chao-Di Chang, Shan-Yun Cheng, Ya-Wen Hung, Chih-Chieh Yang, and Yu-Hsien Chang from Pharmacology Discovery Services Taiwan, Ltd. (PDS), for their outstanding and dedicated efforts in conducting the animal studies, Peter Chi (Institute of Biochemical Sciences, National Taiwan University, Taipei, Taiwan), Chia-Lung Hsieh (Institute of Atomic and Molecular Sciences, Academia Sinica, Taipei, Taiwan) and Sheau-Yann Shieh (Institute of Biomedical Sciences (IBMS), Academia Sinica, Taipei, Taiwan) for their insightful comments and suggestions, and the Core Facilities at IBMS, Academia Sinica (Confocal Microscopy, DNA Sequencing, Flow Cytometry, and Computational Medicine) for their instruments and technical support. Shu-Ping Wang was supported by IBMS, Academia Sinica, and by grants from National Science and Technology Council (NSTC) in Taiwan (MOST110-2628-B-001-017; MOST111-2628-B-001-010; NSTC112-2628-B-001-002). Lan-Hsin Wang was supported by grants from NSTC (MOST107-2311-B-016-001-MY3; MOST110-2311-B-016-002; NSTC112-2311-B-016-001) and the Ministry of National Defense-Medical Affairs Bureau (MND-MAB-D-111086).

## Author Contributions

S.-P.W. conceived the project and wrote the paper. L.-W.Y. performed most of the experiments. S.-P.W. and L.-H.W. supervised the research. Y.-M.C. conducted genome-wide data analyses. L.-H.W. performed fly analyses. J.-W.C. performed cell cycle, colony formation, and apoptosis assays for NCI-H1975 cells. H.-C.H. conducted CUT&RUN analyses. S.W. performed protein interaction assays and histone acid extraction assays. E.P-O., A.E. conducted the Cancer Genome Atlas analysis of expression levels and correlation. L.-W.Y., J.-Y.Y., H.-C.H., W.-M.C. contributed to NCI-H1975 xenograft mouse models. M.-H.K., M.A.E.A., T.-Y.L., E.S. and W.-Y.T. contributed material support and technical assistance.

## References

1. S. Saxena, L. Zou, Hallmarks of DNA replication stress. Mol Cell 82, 2298–2314 (2022).

2. D. Cortez, Replication-Coupled DNA Repair. Mol Cell 74, 866–876 (2019).

3. C. Alabert, A. Groth, Chromatin replication and epigenome maintenance. Nat Rev Mol Cell Biol 13, 153–167 (2012).

4. A. J. Bannister, T. Kouzarides, Regulation of chromatin by histone modifications. Cell Res 21, 381–395 (2011).

5. B. Rondinelli et al., EZH2 promotes degradation of stalled replication forks by recruiting MUS81 through histone H3 trimethylation. Nat Cell Biol 19, 1371–1378 (2017).

6. S. Campbell, I. H. Ismail, L. C. Young, G. G. Poirier, M. J. Hendzel, Polycomb repressive complex 2 contributes to DNA double-strand break repair. Cell Cycle 12, 2675–2683 (2013).

7. K. H. Kim, C. W. Roberts, Targeting EZH2 in cancer. Nat Med 22, 128–134 (2016).

8. A. Piunti, A. Shilatifard, The roles of Polycomb repressive complexes in mammalian development and cancer. Nat Rev Mol Cell Biol 22, 326–345 (2021).

9. A. Piunti, A. Shilatifard, Epigenetic balance of gene expression by Polycomb and COMPASS families. Science (New York, N.Y.) 352, aad9780 (2016).

10. C. Hua et al., KDM6 Demethylases and Their Roles in Human Cancers. Front Oncol 11, 779918 (2021).

11. L. Wang, A. Shilatifard, UTX Mutations in Human Cancer. Cancer Cell 35, 168–176 (2019).

12. T. Miller et al., COMPASS: a complex of proteins associated with a trithorax-related SET domain protein. Proc Natl Acad Sci U S A 98, 12902–12907 (2001).

13. L. H. Wang, M. A. E. Aberin, S. Wu, S. P. Wang, The MLL3/4 H3K4 methyltransferase complex in establishing an active enhancer landscape. Biochem Soc Trans 49, 1041–1054 (2021).

14. F. Tie, R. Banerjee, P. A. Conrad, P. C. Scacheri, P. J. Harte, Histone demethylase UTX and chromatin remodeler BRM bind directly to CBP and modulate acetylation of histone H3 lysine 27. Molecular and cellular biology 32, 2323–2334 (2012).

15. S.-P. Wang et al., A UTX-MLL4-p300 Transcriptional Regulatory Network Coordinately Shapes Active Enhancer Landscapes for Eliciting Transcription. Molecular Cell 67, 308–321 (2017).

16. S. Lee, J. W. Lee, S. K. Lee, UTX, a histone H3-lysine 27 demethylase, acts as a critical switch to activate the cardiac developmental program. Developmental cell 22, 25–37 (2012).

17. G. van Haaften et al., Somatic mutations of the histone H3K27 demethylase gene UTX in human cancer. Nature genetics 41, 521–523 (2009).

18. M. S. Lawrence et al., Discovery and saturation analysis of cancer genes across 21 tumour types. Nature 505, 495–501 (2014).

19. A. Zehir et al., Mutational landscape of metastatic cancer revealed from prospective clinical sequencing of 10,000 patients. Nat Med 23, 703–713 (2017).

20. J. Van der Meulen, F. Speleman, P. Van Vlierberghe, The H3K27me3 demethylase UTX in normal development and disease. Epigenetics 9, 658–668 (2014).

21. W. A. Schulz, A. Lang, J. Koch, A. Greife, The histone demethylase UTX/KDM6A in cancer: Progress and puzzles. Int J Cancer 145, 614–620 (2019).

22. T. L. Lochmann et al., Targeted inhibition of histone H3K27 demethylation is effective in high-risk neuroblastoma. Sci Transl Med 10 (2018).

23. O. A. Romero et al., SMARCA4 deficient tumours are vulnerable to KDM6A/UTX and KDM6B/JMJD3 blockade. Nat Commun 12, 4319 (2021).

24. J. Zhang et al., Targeted inhibition of KDM6 histone demethylases eradicates tumor-initiating cells via enhancer reprogramming in colorectal cancer. Theranostics 10, 10016–10030 (2020).

25. N. Tran, A. Broun, K. Ge, Lysine Demethylase KDM6A in Differentiation, Development, and Cancer. Molecular and cellular biology 40 (2020).

26. J. Abu-Hanna et al., Therapeutic potential of inhibiting histone 3 lysine 27 demethylases: a review of the literature. Clin Epigenetics 14, 98 (2022).

27. N. Cancer Genome Atlas Research et al., The Cancer Genome Atlas Pan-Cancer analysis project. Nature genetics 45, 1113–1120 (2013).

28. Y. Wei, C. A. Mizzen, R. G. Cook, M. A. Gorovsky, C. D. Allis, Phosphorylation of histone H3 at serine 10 is correlated with chromosome condensation during mitosis and meiosis in Tetrahymena. Proc Natl Acad Sci U S A 95, 7480–7484 (1998).

29. A. Van Hooser, D. W. Goodrich, C. D. Allis, B. R. Brinkley, M. A. Mancini, Histone H3 phosphorylation is required for the initiation, but not maintenance, of mammalian chromosome condensation. J Cell Sci 111 (**Pt 23**), 3497–3506 (1998).

30. C. Gelot, I. Magdalou, B. S. Lopez, Replication stress in Mammalian cells and its consequences for mitosis. Genes (Basel) 6, 267–298 (2015).

31. J. Guirouilh-Barbat, S. Lambert, P. Bertrand, B. S. Lopez, Is homologous recombination really an error-free process? Front Genet 5, 175 (2014).

32. A. Ray Chaudhuri, A. Nussenzweig, The multifaceted roles of PARP1 in DNA repair and chromatin remodelling. Nat Rev Mol Cell Biol 18, 610–621 (2017).

33. B. A. Edgar, T. L. Orr-Weaver, Endoreplication cell cycles: more for less. Cell 105, 297–306 (2001).

34. H. O. Lee, J. M. Davidson, R. J. Duronio, Endoreplication: polyploidy with purpose. Genes Dev 23, 2461–2477 (2009).

35. M. P. Hammond, C. D. Laird, Control of DNA replication and spatial distribution of defined DNA sequences in salivary gland cells of Drosophila melanogaster. Chromosoma 91, 279–286 (1985).

36. P. J. Follette, R. J. Duronio, P. H. O’Farrell, Fluctuations in cyclin E levels are required for multiple rounds of endocycle S phase in Drosophila. Curr Biol 8, 235–238 (1998).

37. A. Weiss, A. Herzig, H. Jacobs, C. F. Lehner, Continuous Cyclin E expression inhibits progression through endoreduplication cycles in Drosophila. Curr Biol 8, 239–242 (1998).

38. N. Zielke et al., Control of Drosophila endocycles by E2F and CRL4(CDT2). Nature 480, 123–127 (2011).

39. B. K. Tripathi, K. D. Irvine, The wing imaginal disc. Genetics 220 (2022).

40. E. Petermann, M. L. Orta, N. Issaeva, N. Schultz, T. Helleday, Hydroxyurea-stalled replication forks become progressively inactivated and require two different RAD51-mediated pathways for restart and repair. Mol Cell 37, 492–502 (2010).

41. J. Du et al., Quantitative assessment of HR and NHEJ activities via CRISPR/Cas9-induced oligodeoxynucleotide-mediated DSB repair. DNA Repair (Amst) 70, 67–71 (2018).

42. J. H. Vissers, M. van Lohuizen, E. Citterio, The emerging role of Polycomb repressors in the response to DNA damage. J Cell Sci 125, 3939–3948 (2012).

43. D. M. Chou et al., A chromatin localization screen reveals poly (ADP ribose)-regulated recruitment of the repressive polycomb and NuRD complexes to sites of DNA damage. Proc Natl Acad Sci U S A 107, 18475–18480 (2010).

44. A. Chang et al., Recruitment of KMT2C/MLL3 to DNA Damage Sites Mediates DNA Damage Responses and Regulates PARP Inhibitor Sensitivity in Cancer. Cancer Res 81, 3358–3373 (2021).

45. A. M. Weber, A. J. Ryan, ATM and ATR as therapeutic targets in cancer. Pharmacol Ther 149, 124–138 (2015).

46. L. Mei, J. Zhang, K. He, J. Zhang, Ataxia telangiectasia and Rad3-related inhibitors and cancer therapy: where we stand. J Hematol Oncol 12, 43 (2019).

47. B. Shi et al., UTX condensation underlies its tumour-suppressive activity. Nature 597, 726–731 (2021).

48. F. Michelini et al., Damage-induced lncRNAs control the DNA damage response through interaction with DDRNAs at individual double-strand breaks. Nat Cell Biol 19, 1400–1411 (2017).

49. S. Francia et al., Site-specific DICER and DROSHA RNA products control the DNA-damage response. Nature 488, 231–235 (2012).

50. W. Wei et al., A role for small RNAs in DNA double-strand break repair. Cell 149, 101–112 (2012).

51. J. F. Haince et al., PARP1-dependent kinetics of recruitment of MRE11 and NBS1 proteins to multiple DNA damage sites. The Journal of biological chemistry 283, 1197–1208 (2008).

52. A. Aguilera, T. Garcia-Muse, Causes of genome instability. Annu Rev Genet 47, 1–32 (2013).

53. J. H. Kim et al., UTX and MLL4 coordinately regulate transcriptional programs for cell proliferation and invasiveness in breast cancer cells. Cancer Res 74, 1705–1717 (2014).

54. G. Xie et al., UTX promotes hormonally responsive breast carcinogenesis through feed-forward transcription regulation with estrogen receptor. Oncogene 36, 5497–5511 (2017).

55. J. H. Taube et al., The H3K27me3-demethylase KDM6A is suppressed in breast cancer stem-like cells, and enables the resolution of bivalency during the mesenchymal-epithelial transition. Oncotarget 8, 65548–65565 (2017).

56. N. Yan et al., GSKJ4, an H3K27me3 demethylase inhibitor, effectively suppresses the breast cancer stem cells. Exp Cell Res 359, 405–414 (2017).

57. N. Dalpatraj, A. Naik, N. Thakur, GSK-J4: An H3K27 histone demethylase inhibitor, as a potential anti-cancer agent. Int J Cancer 153, 1130–1138 (2023).

58. H. Katagi et al., Radiosensitization by Histone H3 Demethylase Inhibition in Diffuse Intrinsic Pontine Glioma. Clin Cancer Res 25, 5572–5583 (2019).

59. S. B. Frank, V. V. Schulz, C. K. Miranti, A streamlined method for the design and cloning of shRNAs into an optimized Dox-inducible lentiviral vector. BMC Biotechnol 17, 24 (2017).

60. R. X. Tsai et al., TERRA regulates DNA G-quadruplex formation and ATRX recruitment to chromatin. Nucleic Acids Res 50, 12217–12234 (2022).

