## Supplemental Information for "The tumor-maintaining function of UTX/KDM6A in DNA replication and the PARP1-dependent repair pathway"

1. Program in Molecular Medicine, National Yang-Ming University and Academia Sinica, Taipei, Taiwan; 2. Institute of Biomedical Sciences, Academia Sinica, Taipei, Taiwan; 3. Program for Cancer Biology and Drug Discovery, China Medical University, Taichung, Taiwan 4. Taiwan International Graduate Program in Molecular Medicine, Academia Sinica and National Yang Ming Chiao Tung University, Taipei, Taiwan; 5. Taiwan International Graduate Program in Molecular and Cellular Biology, Academia Sinica and Graduate Institute of Life Science, National Defense Medical Center, Taipei, Taiwan; 6. Computational Genomics Division, National Institute of Genomic Medicine, Mexico City, Mexico; 7. Department and Graduate Institute of Biochemistry, National Defense Medical Center, Taipei, Taiwan.

*Shu-Ping Wang.

**This PDF file includes:**

Supporting text
SI References
Figures S1 to S10
Tables S1 to S3

#### Supporting Information Text

#### Extended Materials and Methods

##### RNA Sequencing (RNA-seq) and Data Analysis

For RNA-seq analysis, RNA was extracted using the RNAspin Mini Kit (Cytiva, Cat#25050071). Subsequently, sequencing libraries were constructed with the TruSeq Stranded mRNA Library Prep Kit (Illumina) following the manufacturer's instructions. The resulting libraries were enriched via PCR and purified using the AMPure XP system (Beckman Coulter). Library quality was assessed using the Qsep400 System (Bioptic Inc.), and quantification was performed using the Qubit 2.0 Fluorometer (Thermo Fisher Sc.). The qualified libraries were sequenced on an Illumina NovaSeq 6000 platform, generating 150 bp paired-end reads by Genomics BioSci. & Tech. For RNA-seq data analysis, raw reads underwent preprocessing with Trimmomatic (1) to eliminate adapters and low-quality sequences (cutoff = Q20). Processed reads were tehn aligned to the human genome (GRCh38) using Hisat2 (2). The expression levels (read count and TPM) of each gene were quatified using Stringtie (2) and human gene annotation from Ensembl (release 104). Differentially expressed genes (DEGs) were identified using the DESeq2 R package. KEGG pathway enrichment test was conducted using WebGestalt (3) with FDR < 0.05. Additionally, Gene Set Enrichment Analysis (GSEA) was performed using GSEA v4.2.3 software (4) with 1,000 permutations.

##### CUT&RUN

CUT&RUN assays were conducted in accordance with the protocol provided by Cell Signaling Technology for their CUT & RUN Assay Kit (Cat#91931). Briefly, 250,000 cells were harvested and lysed using DNA extraction buffer, followed by two washes with 1mL of wash buffer per sample. Subsequently, the cell mixture was combined with Concanavalin A beads to activate and capture cells. The histone H3K27me3 antibody (anti-H3K27me3; Abcam, Cat#ab1791) were employed to penetrate the cells, and the samples were then incubated overnight at 4°C on a rotating platform. MNase digestion was initiated with 2 mM $CaCl_2$ and carried out for 30 min at 4°C with the beads. Following this, Stop buffer containing spike-in DNA was added to each sample and incubated at 37°C for 30 min without shaking to release DNA fragments into solution. Libraries were prepared using the ChIP-seq DNA Library Prep Kit (Illumina, Cat#56795) with 13 cycles of amplification. Purification of the libraries was performed using AMPure XP beads (Beckman Coulter, #A63880), and sequencing was carried out on an Illumina Novaseq X Plus platform using 150PE reads.

##### Analysis of expression levels and correlation

The correlation analysis was performed using the dataset from The Cancer Genome Atlas (TCGA) Pan-Cancer analysis project (5). We utilized a dataset comprising 11,069 samples. From this dataset, we specifically extracted 217 samples derived from lung tissue with neoplastic conditions due to lung cancer exhibited the most notable correlations in our preliminary study. Our analysis relied on an expression matrix encompassing bulk mRNA expression levels, which were harmonized using the ComBat algorithm (6). This preprocessing step ensured consistency and minimized batch effects across samples. To explore the simultaneous correlation between the genes of interest (UTX/KDM6A and MLL4/KMT2D) and genes associated with DNA replication, we employed 3D scatterplots. In these plots, KDM6A and KMT2D were represented on the x and y axes, while the functional gene associated with DNA duplication and replication was plotted on the y-axis, respectively, while the functional gene associated with DNA replication was depicted on the z-axis. To quantify the linear correlation and assess the goodness-of-fit, we utilized multivariate least squares regression analysis. This statistical approach enabled us to estimate regression coefficients and determine the coefficient of determination (R^2). The p-values for the coefficients were calculated by adjusting for a t-distribution against the null hypothesis of being zero. Additionally, 2D scatterplots were generated with KMT2D or KDM6A on the x-axis and the functional gene associated with DNA replication on the y-axis to identify direct linear correlations, considering the regression coefficient size enhancement provided by multivariate least squares regression.

#### SI References

1. A. M. Bolger, M. Lohse, B. Usadel, Trimmomatic: a flexible trimmer for Illumina sequence data. *Bioinformatics* **30**, 2114-2120 (2014).
2. M. Pertea, D. Kim, G. M. Pertea, J. T. Leek, S. L. Salzberg, Transcript-level expression analysis of RNA-seq experiments with HISAT, StringTie and Ballgown. *Nat Protoc* **11**, 1650-1667 (2016).
3. Y. Liao, J. Wang, E. J. Jaehnig, Z. Shi, B. Zhang, WebGestalt 2019: gene set analysis toolkit with revamped UIs and APIs. *Nucleic Acids Res* **47**, W199-W205 (2019).
4. A. Subramanian *et al.*, Gene set enrichment analysis: a knowledge-based approach for interpreting genome-wide expression profiles. *Proc Natl Acad Sci U S A* **102**, 15545-15550 (2005).
5. N. Cancer Genome Atlas Research *et al.*, The Cancer Genome Atlas Pan-Cancer analysis project. *Nature genetics* **45**, 1113-1120 (2013).
6. Y. Zhang, G. Parmigiani, W. E. Johnson, ComBat-seq: batch effect adjustment for RNA-seq count data. *NAR Genom Bioinform* **2**, lqaa078 (2020).

### Supplementary Figure S1

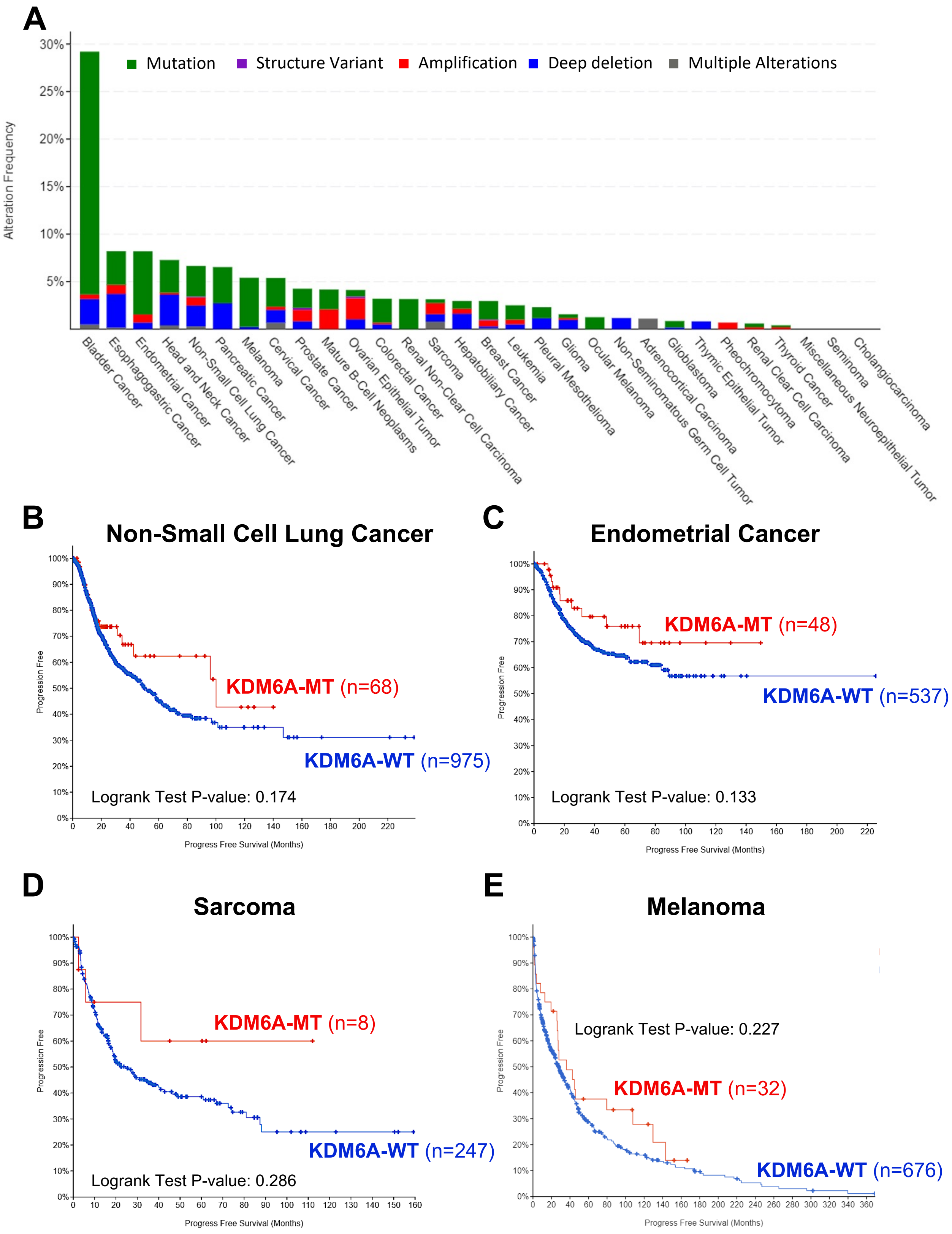

**Figure S1. General correlation between UTX/KDM6A mutation states and cancer patients' clinical outcomes.**

(A) KDM6A mutation frequencies across various human cancer types as obtained from cBioPortal. (B–E) Kaplan-Meier survival curves of progression-free survival (PFS) in non-small cell lung cancer (B), endometrial cancer (C), sarcoma (D), or melanoma (E) patients with WT and MT forms of KDM6A from the cBioPortal dataset.

### Supplementary Figure S2

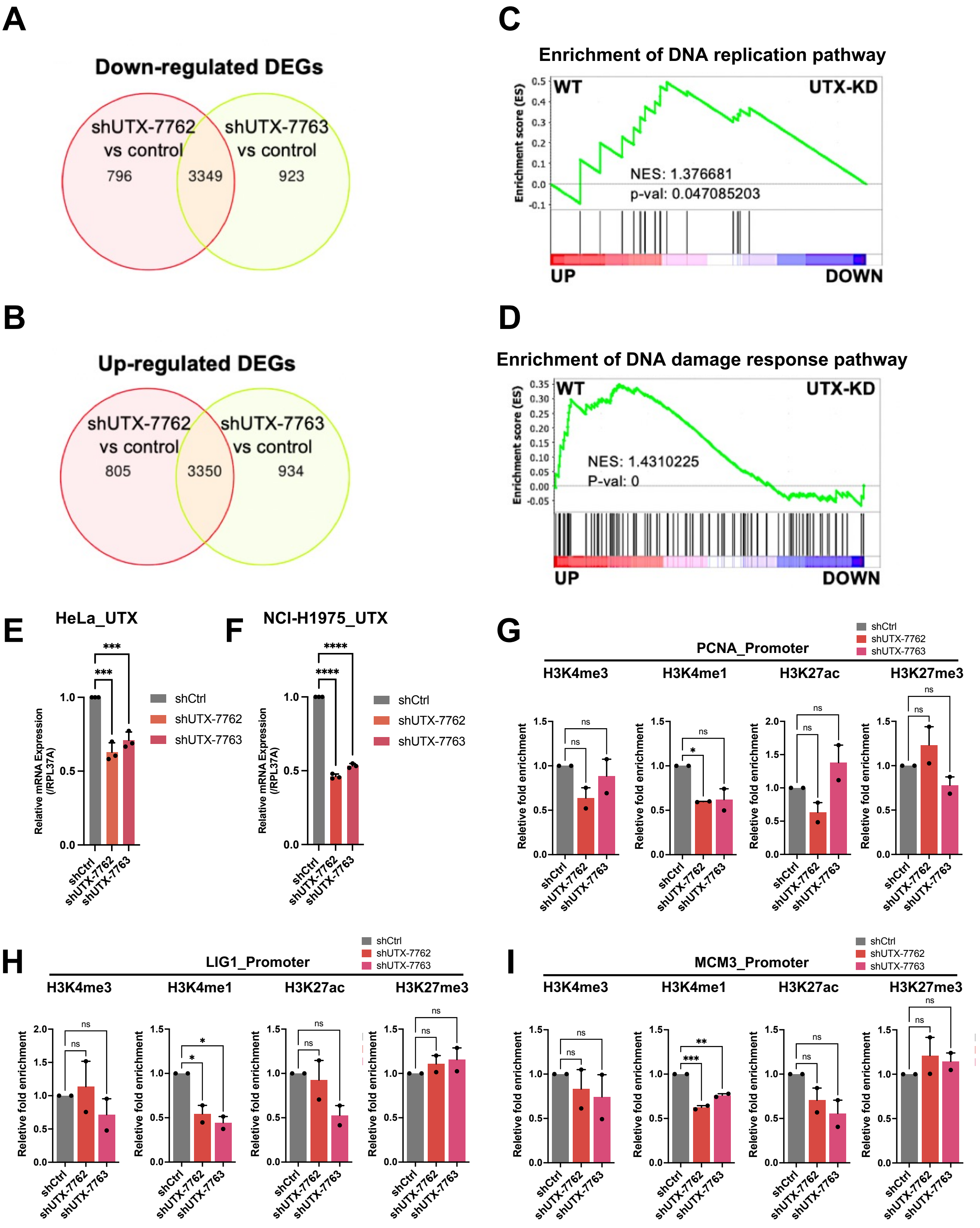

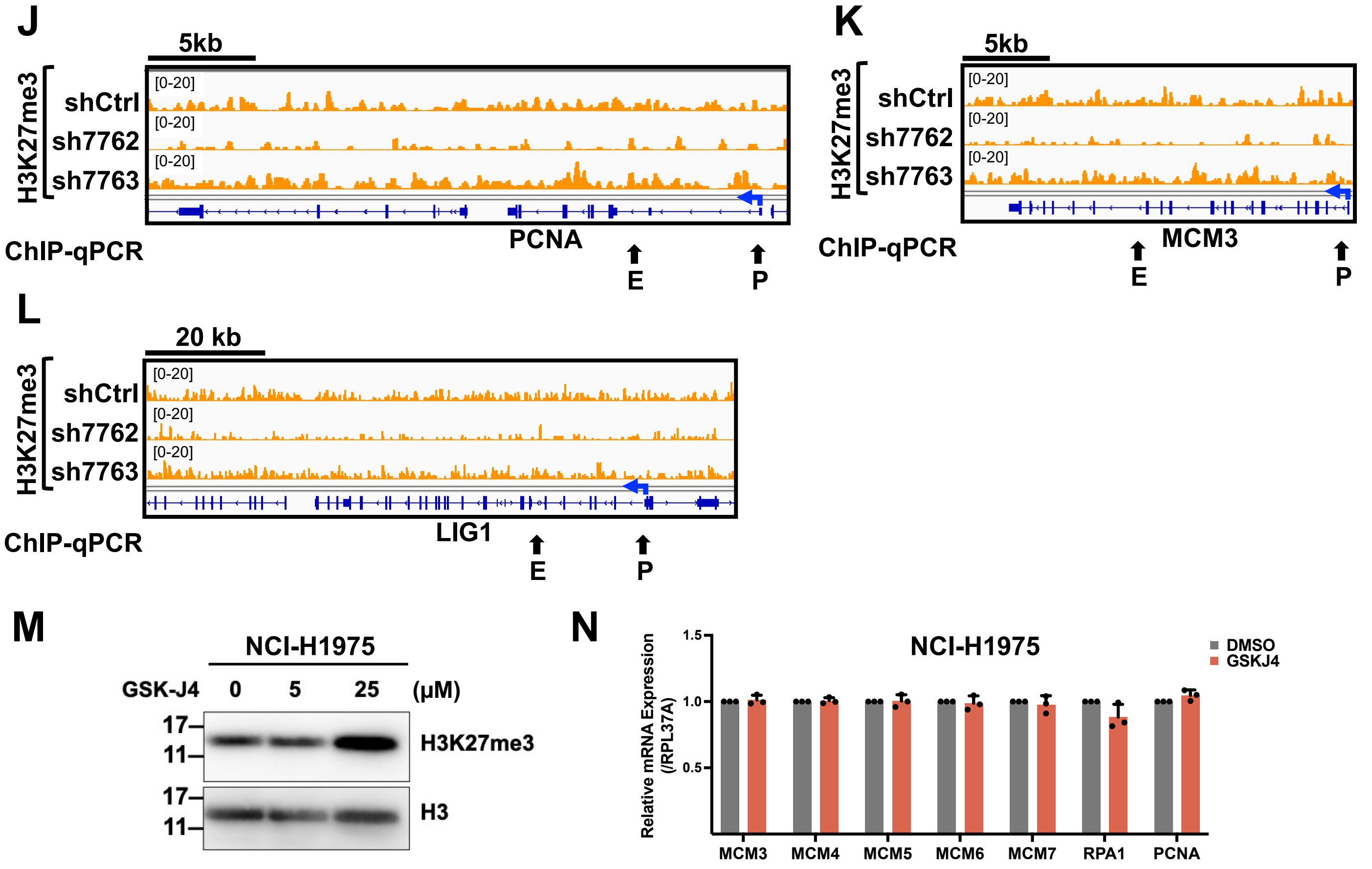

**Figure S2. The potential involvement of UTX/KDM6A in regulating DNA replication and repair pathways.**

(A and B) Venn diagrams illustrate differentially-expressed genes (DEGs) that were either down- or up-regulated by UTX/KDM6A depletion (shUTX-7762 or shUTX-7763) compared with control (shCtrl) HeLa cells. (C, D) Enrichment score plots from Gene-Set Enrichment Analysis (GSEA) demonstrate significant downregulation of genes involved in DNA replication (C) or DNA damage response (D) in UTX/KDM6A-knockdown HeLa cells compared to control cells. Up and down indicate the relative gene up- or downregulation post UTX/KDM6A depletion. The analysis employed GO_BP. NES, normalized enrichment score; p-val, p-value. (E and F) RT-qPCR analysis of UTX/KDM6A mRNA levels in control (shCtrl) and UTX/KDM6A-shRNAs (shUTX-7762 or shUTX-7763)-transduced HeLa (E) or NCI-H1975 (F) cells. ***p < 0.001, ****p < 0.0001 (Ordinary one-way ANOVA). (G–I) Enrichments of histone marks (H3K4me3, H3K4me1, H3K27ac, and H3K27me3) at three representative gene promoters in control (shCtrl) and UTX/KDM6A-shRNAs (shUTX-7762 or shUTX-7763)-transduced NCI-H1975 cells. (G) *PNCA*. (H) *LIG1*. (I) *MCM3*. Fold enrichment values were normalized to H3. Relative fold enrichment was further normalized to control cells (n = 2 biological replicates from 2 independent experiments). *p < 0.05, **p < 0.01, ***p < 0.001, ns, no significant (Ordinary one-way ANOVA). (J–L) Screen shot of CUT&RUN profiles for H3K27me3 at selected gene loci in control (shCtrl) and UTX/KDM6A-shRNAs (shUTX-7762 or shUTX-7763)-transduced NCI-H1975 cells. (J) *PNCA*. (K) *LIG1*. (L) *MCM3*. Arrows indicate the corresponding genomic locations of ChIP-qPCR amplicons. E, enhancer; P, promoter. (M) Levels of H3K27me3 in histone extracts from NCI-H1975 cells treated with GSK-J4 or left untreated, assessed by Western blot analysis. Histone extracts were prepared by acid extraction. Histone H3 served as a loading control. (N) RT-qPCR analysis of mRNA levels of DNA replication-associated genes conducted in NCI-H1975 cells treated with GSK-J4 or left untreated. RPL37A was used as an endogenous control. Data are presented as mean ± SEM from three independent experiments (Two-way ANOVA).

### Supplementary Figure S3

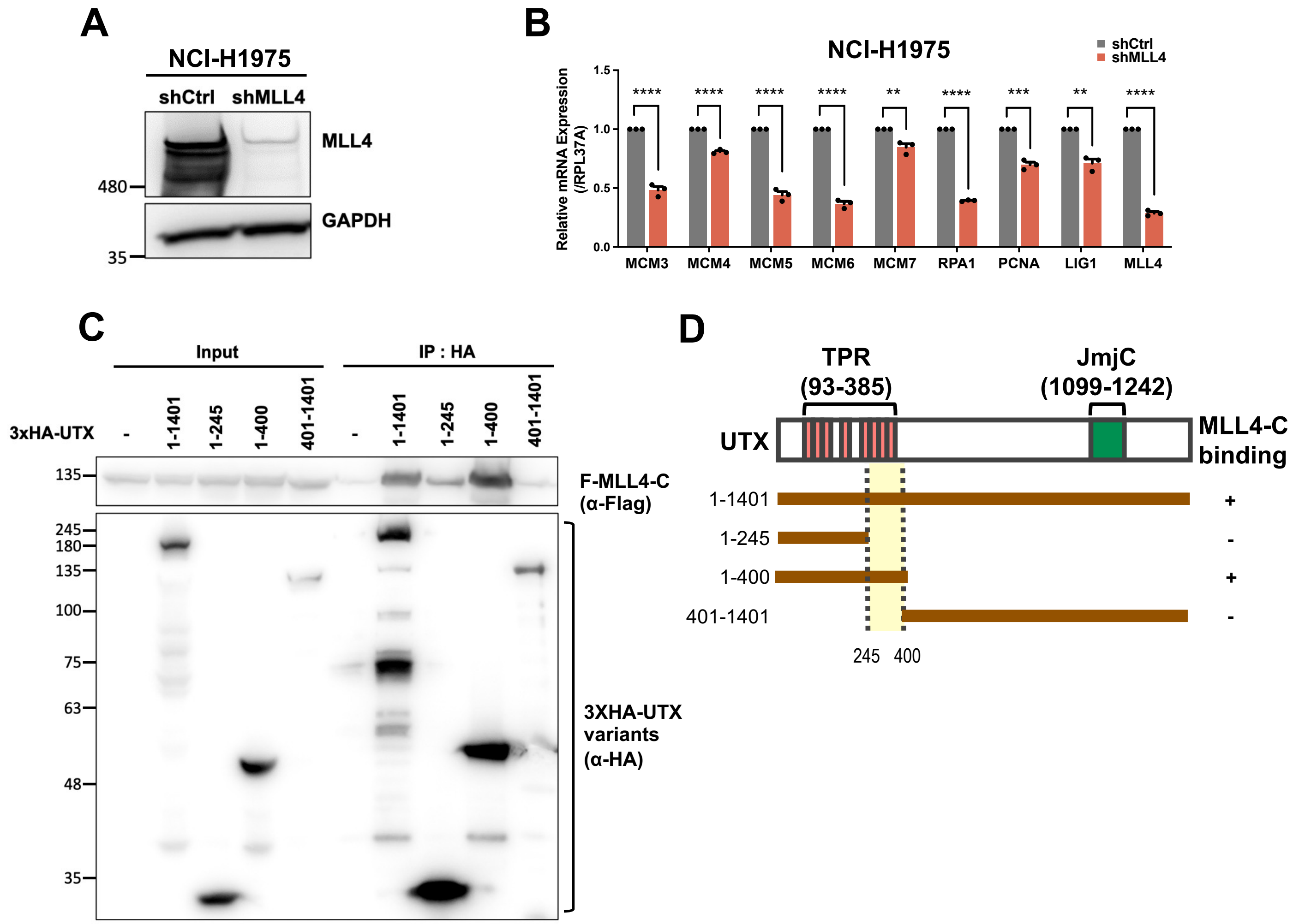

**Figure S3. The interplay between UTX/KDM6A and MLL4/KMT2D is pivotal for the activation of DNA replication-associated genes.**
(A) Western blot analysis with the specified antibodies of lysates obtained from control (shCtrl) and MLL4/KMT2D-shRNA (shMLL4)-transduced NCI-H1975 cells. (B) RT-qPCR analysis of mRNA levels of DNA replication-associated genes conducted in NCI-H1975 cells with control (shCtrl) or MLL4/KMT2D-shRNA (shMLL4). RPL37A served as an endogenous control. Data are presented as mean ± SEM from three independent experiments. $^{**}p < 0.01$, $^{***}p < 0.001$, $^{****}p < 0.0001$ (Two-way ANOVA). (C) Co-immunoprecipitation of Flag-tagged MLL4-C (C-terminal region, amino acids 4507-5537) and different 3XHA-tagged UTX deletion variants from HEK293 whole cell extract following incubation with anti-HA antibody. Immunoprecipitates were subjected to immunoblotting with the specified antibodies. (D) Schematic illustration and summary outlining the interaction between MLL4-C and different deletion fragments of UTX/KDM6A.

### Supplementary Figure S4

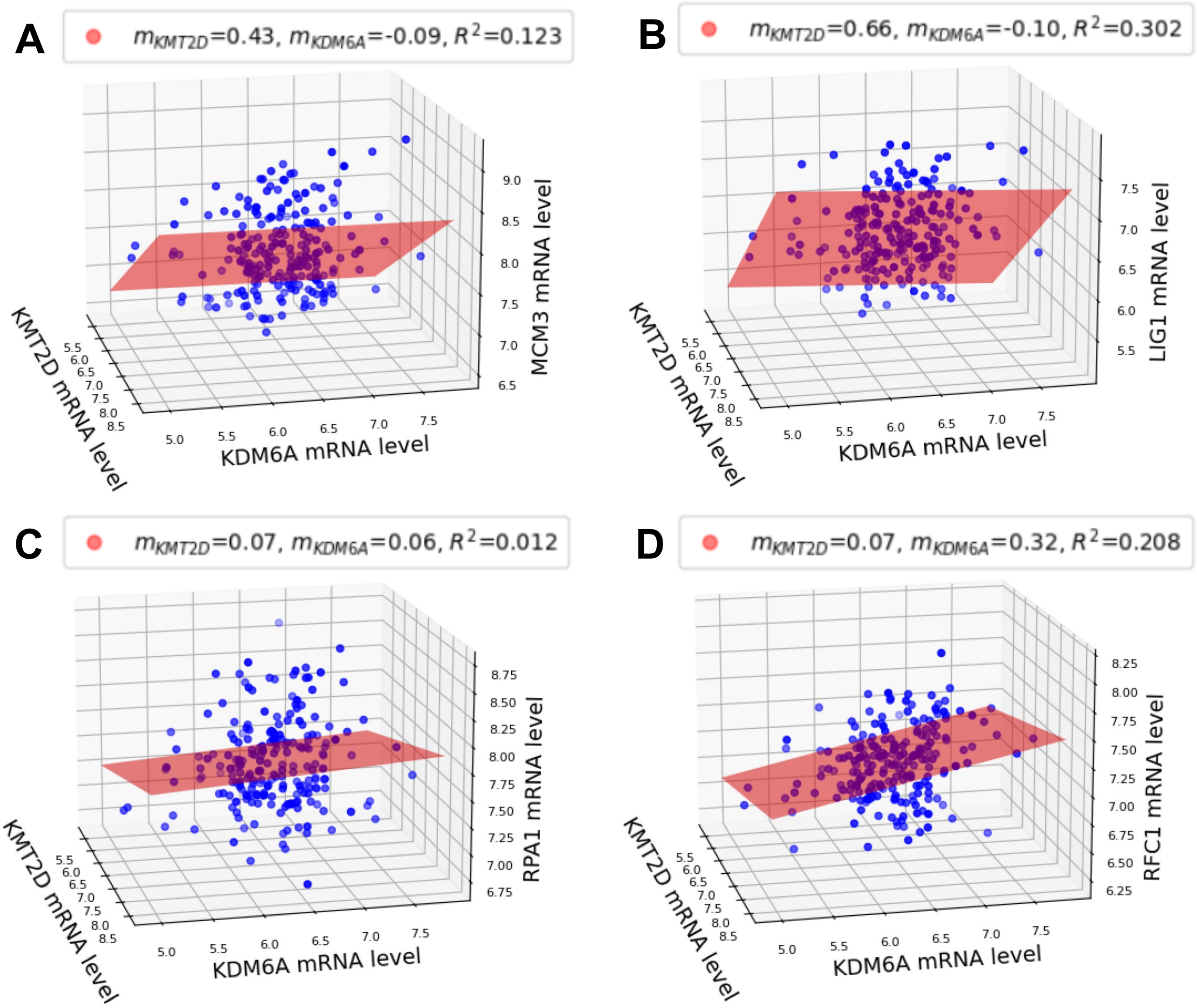

**Figure S4. Correlation between UTX/KDM6A, MLL4/KMT2D, and DNA replication-associated genes in NSCLC.**
(**A**) MCM3: $p_{KMT2D}$ = 8.95E-01, $p_{KDM6A}$ = 7.02E-10, (**B**) LIG1: $p_{KMT2D}$ = 3.81E-01, $p_{KDM6A}$ = 1.86E-04, (**C**) RPA1: $p_{KMT2D}$ = 3.13E-01, $p_{KDM6A}$ = 1.01E-10, (**D**) RFC1: $p_{KMT2D}$ = 1.39E-04, $p_{KDM6A}$ = 1.02E-10.

### Supplementary Figure S5

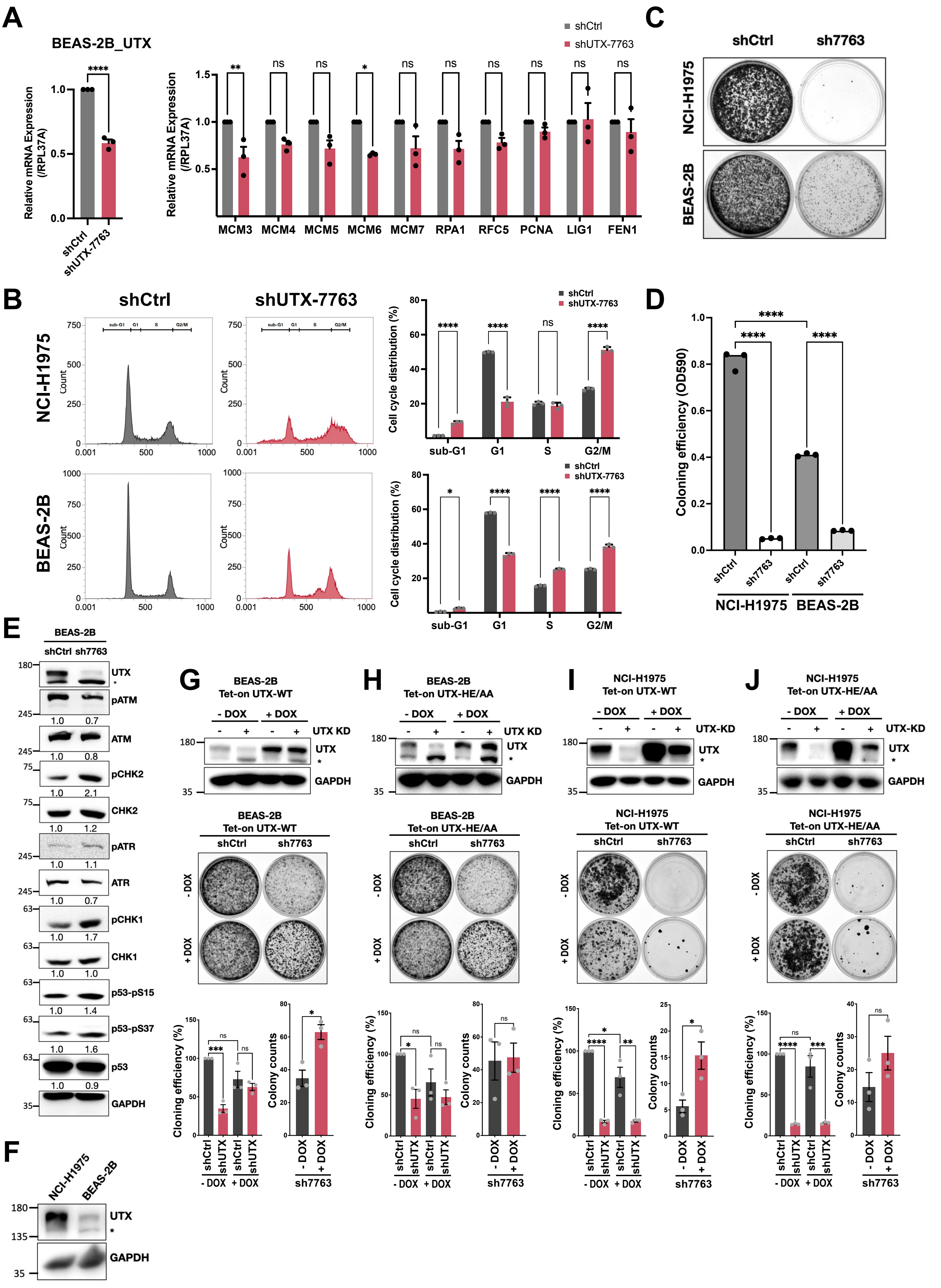

**Figure S5. The limited impact of UTX/KDM6A loss on normal cells.**
(A) Left: RT-qPCR analysis depicts UTX/KDM6A mRNA levels in control (shCtrl) and UTX-shRNA (shUTX-7763)-transduced BEAS-2B cells. Right: RT-qPCR analysis depicts mRNA levels of DNA replication-associated genes in BEAS-2B cells with control or UTX-shRNA (shUTX-7763), with RPL37A as an endogenous control. Data represent mean ± SEM from three independent experiments. ns, no significant; *$p < 0.05$, **$p < 0.01$ (Two-way ANOVA). (B) Left: Flow cytometric analysis of cell cycle phases for Dox-inducible (Tet-on) shUTX-expressing NCI-H1975, BEAS-2B, or respective control (shCtrl) cells with or without Dox treatment. Sub-G1, G1, S, and G2/M phases were calculated from the PI-labeled histograms after 72-h treatment with 50 ng/mL Dox. Right: Quantification of cells in each phase is presented as mean ± SEM for three independent experiments. (C) Stable UTX-knockdown or control BEAS-2B cells were subjected to colony formation assays, with representative plate images shown. (D) Colony formation ability of stable cell clones in (C) determined by crystal violet assays. Quantification is presented as mean ± SEM for three independent experiments. ****$p < 0.0001$ (Ordinary one-way ANOVA). (E) Western blot analysis with indicated antibodies of lysates from control (shCtrl) and UTX/KDM6A-shRNA (shUTX-7763)-transduced BEAS-2B cells. Densitometric values normalized to GAPDH are indicated below the immunoblot panels. * a non-specific band. (F) Western blot analysis with indicated antibodies of lysates from NCI-H1975 and BEAS-2B cells. * a non-specific band. (G–J) Growth inhibitory effects of UTX/KDM6A-knockdown on Dox-induced re-expression of wild-type (WT) or a demethylase-dead mutant (HE/AA) of UTX/KDM6A in BEAS-2B and NCI-H1975 cells. Top: Western blot analysis of lysates from control (shCtrl) and shUTX-7763 (sh7763)-transduced BEAS-2B and NCI-H1975 cells expressing Dox-inducible (Tet-on) WT or HE/AA UTX/KDM6A with or without Dox treatment. Stable cell clones with different conditions were subjected to colony formation assays, with representative plate images shown in the middle. Bottom: Quantification presented as mean ± SEM for three independent experiments.

#### Supplementary Figure S6

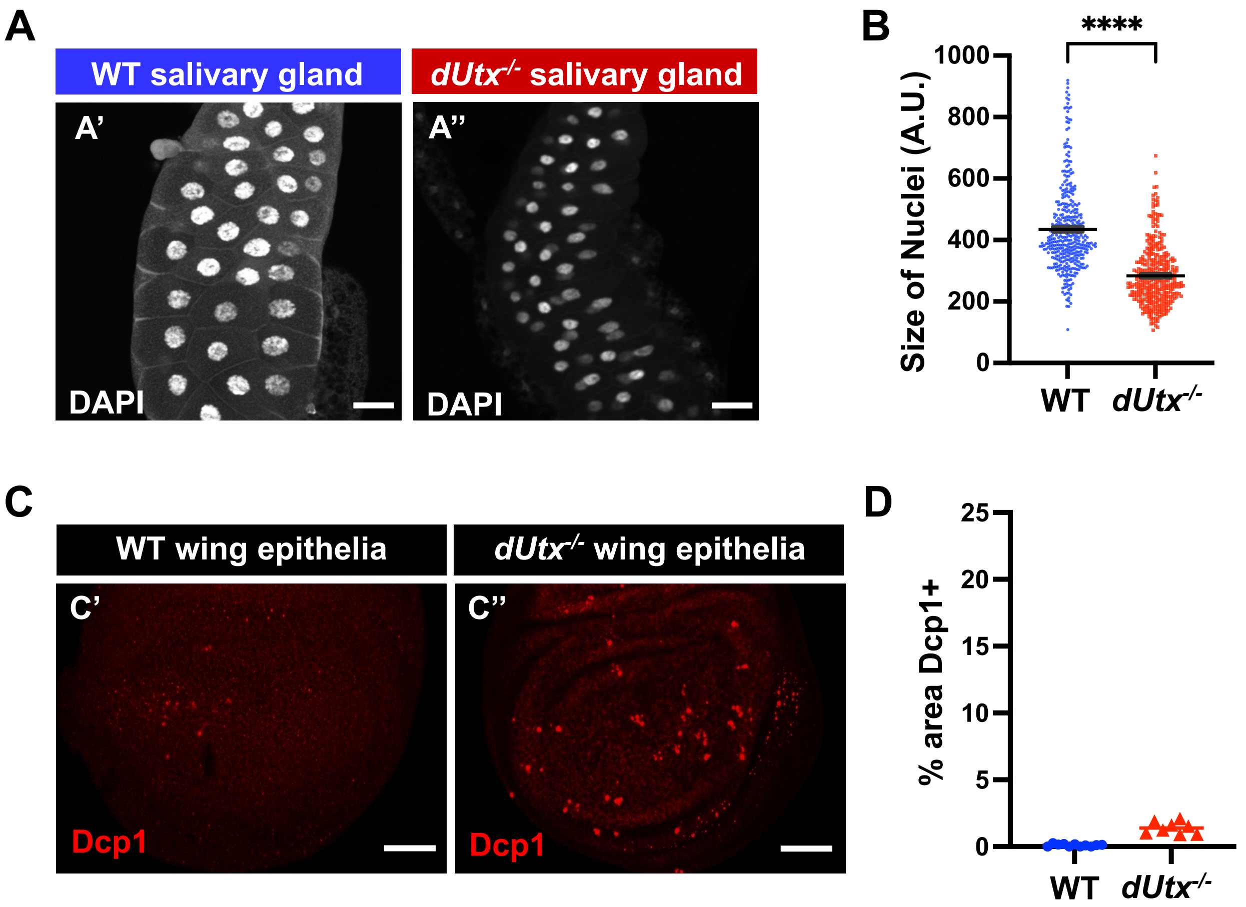

**Figure S6. The impact of UTX/KDM6A loss on normal tissues is relatively minor.**
(A) Salivary glands of *Drosophila* wandering larvae from wild-type (WT, A') or dUtx homozygous mutant (dUTX$^{-/-}$, A'') were stained with DAPI, respectively. (B) Quantification data of nuclear size measurements in salivary glands of WT (B') or dUTX$^{-/-}$ (B'') genotypes. Nuclei were calibrated using the Cellpose algorithm followed by ImageJ measurements for individual nuclear size. A.U. represents arbitrary unit. Each dot represents one nucleus. Data are presented as mean ± SEM (n=15 salivary glands for WT; n=7 salivary glands for dUTX$^{-/-}$). Scale bar: 50 μm. ****P value <0.0001 (Unpaired t-test). (C) *Drosophila* wing epithelial tissue, specifically the pouch region of the third-instar larval wing imaginal discs, assayed for apoptosis by anti-Dcp1 staining (red). WT (C') or dUTX$^{-/-}$ (C'') genotypes are shown respectively. (D) The percentage of Dcp1-positive staining in the pouch region from (C') and (C'') was quantified. Each dot represents one wing disc. WT: 0.1025%, dUtx$^{-/-}$ : 1.419%. Data are presented as mean ± SEM (n=11 wing epithelia for WT; n=8 wing epithelia for dUTX$^{-/-}$). Scale bar: 50 μm. Error bars indicate the SEM of the results of indicated genotypes. ****P value<0.0001 (Unpaired t-test).

### Supplementary Figure S7

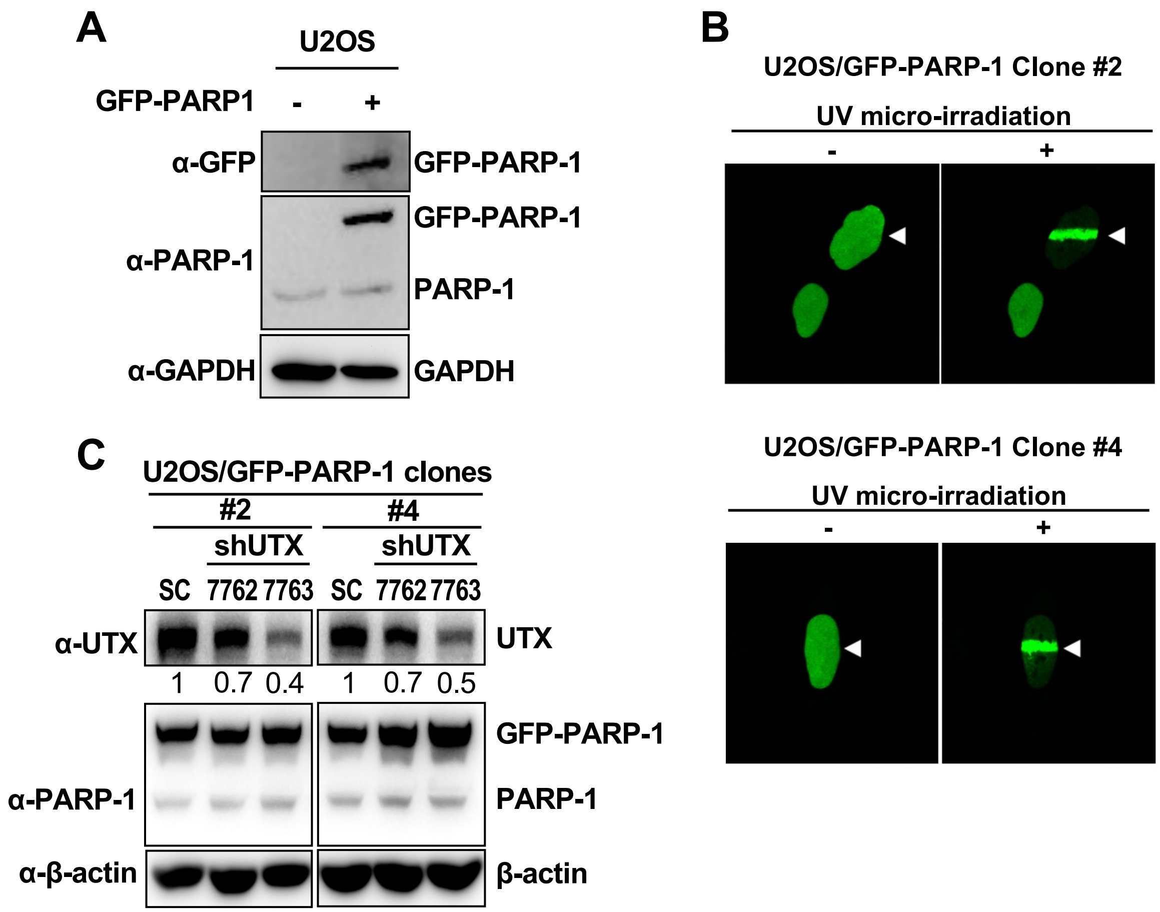

**Figure S7. Establishing a laser-induced DNA damage system to monitor DNA damage-induced PARP-1 recruitment.**
(A) Western blot analysis using the indicated antibodies of lysates from U2OS cells stably transfected with either the empty vector or the expression construct encoding GFP-tagged PARP-1 (GFP-PARP-1). (B) UV laser micro-irradiation was performed on independently stably transfected subclones expressing GFP-PARP-1 (clones #2 and #4), followed by analysis of GFP-PARP-1 recruitment to the laser-induced DNA damage sites using confocal microscopy. (C) Western blot analysis using the indicated antibodies of lysates from two independent U2OS subclones expressing GFP-PARP-1 (U2OS/GFP-PARP-1 #2 and #4) with stable transduction of shCtrl, shUTX-7762, or shUTX-7763. Ctrl, control.

### Supplementary Figure S8

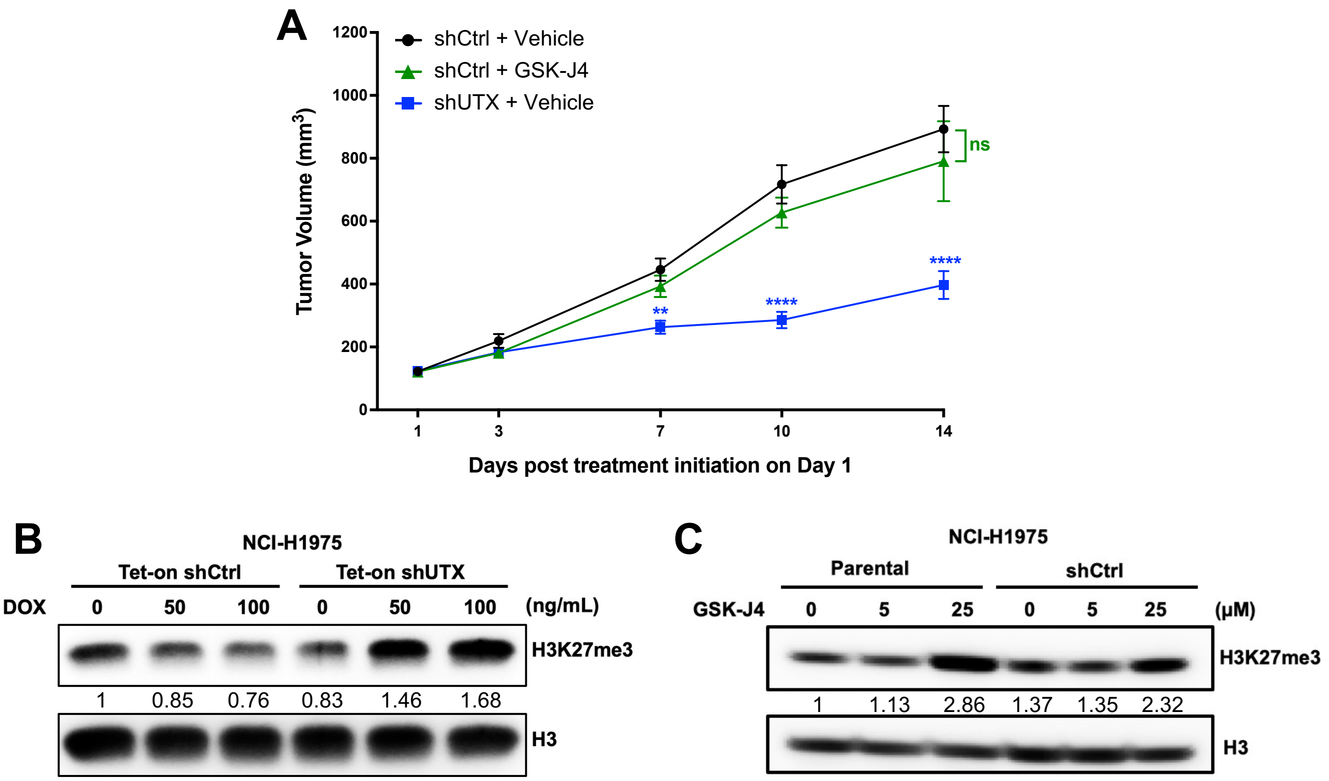

**Figure S8. GSK-J4 does not significantly affect the growth of NCI-H1975 tumors.** (A) Tumor growth rates of Dox-inducible (Tet-on) shUTX-expressing NCI-H1975 or control (shCtrl) cells upon treatment with vehicle, Dox, or GSK-J4 as indicated in xenograft assays (n=8). Data represent means ± SEM. ns, no significant; **$p < 0.01$, ****$p < 0.0001$ (Two-way ANOVA followed by Bonferroni correction). (B) Levels of H3K27me3 in histone extracts from Dox-induced (Tet-on) UTX/KDM6A-knockdown (shUTX) NCI-H1975 or control (shCtrl) cells, as assessed by Western blot analysis. (C) Levels of H3K27me3 in histone extracts from NCI-H1975 (parental or shCtrl) cells treated with GSK-J4 or left untreated, as assessed by Western blot analysis. Histone extracts were obtained through acid extraction. Histone H3 served as a loading control. Numbers below the immunoblot panels indicate the densitometric values normalized to the respective H3 value. Ctrl, control. Dox, doxycycline.

### Supplementary Figure S9

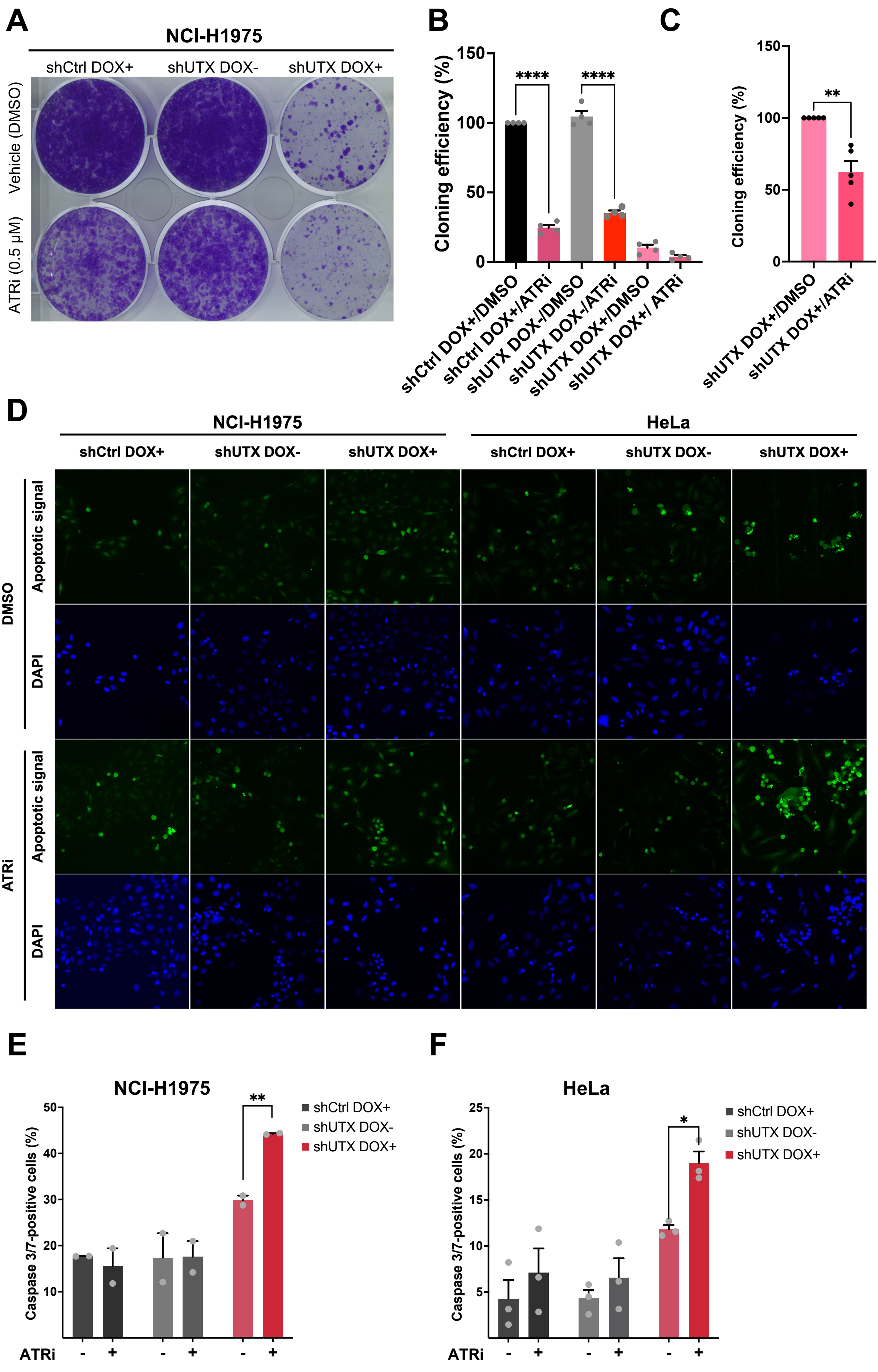

**G**

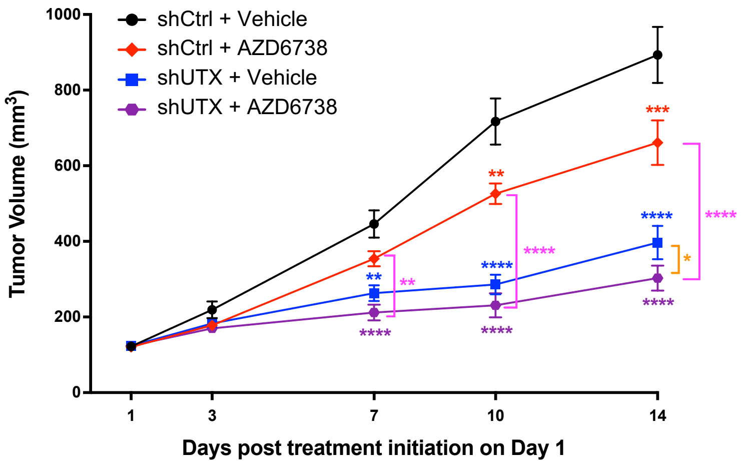

**Figure S9. The combined inhibition of UTX/KDM6A and ATR demonstrates synergistic anti-tumor effects.**

(A) Dox-inducible (Tet-on) shUTX-expressing NCI-H1975 or control (shCtrl) cells were treated with vehicle (DMSO) or ATR inhibitor (ATRi, AZD6738, 0.5 μM) with or without Dox (50 ng/ml), followed by colony formation assays. Representative images of the plates are shown. (B) The colony formation ability of each combination treatment in (A) was assessed by crystal violet assays. Quantification is presented as mean ± SEM for three independent experiments. ****$p < 0.0001$ (Ordinary one-way ANOVA). (C) The colony formation ability of single inhibition of UTX/KDM6A or dual inhibition of UTX/KDM6A and ATR was evaluated by colony numbers using ImageJ measurements. Quantification is shown as mean ± SEM for three independent experiments. **$p < 0.01$ (Ordinary one-way ANOVA). (D) Representative images of Dox-inducible (Tet-on) shUTX-expressing NCI-H1975, HeLa or the respective control (shCtrl) cells treated with Dox (50 ng/ml), ATRi (AZD6738, 0.5 μM for NCI-H1975; 0.625 μM for HeLa) or the indicated combinations in the presence of a green fluorescent caspase-3/7 activatable dye. (E and F) The proportion of cells with caspase-3/7 staining in NCI-H1975 (E) and HeLa (F) was quantified using ImageJ measurements. Quantification is presented as mean ± SEM for two or three independent experiments. *$p < 0.05$, **$p < 0.01$ (Two-way ANOVA). (G) Tumor growth rates of Dox-inducible (Tet-on) shUTX-expressing NCI-H1975 or control (shCtrl) cells upon treatment with vehicle, Dox, or AZD6738 as indicated in xenograft assays (n=8). Data represented means ± SEM. *$p < 0.05$, **$p < 0.01$, ***$p < 0.001$, ****$p < 0.0001$ (Two-way ANOVA followed by Bonferroni correction).

### Supplementary Figure S10

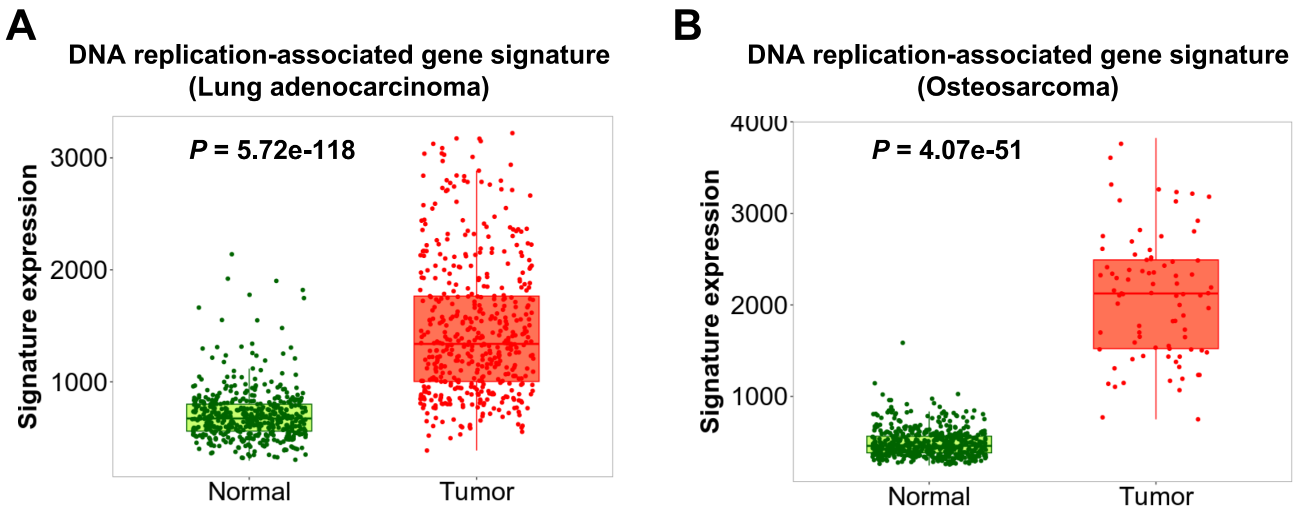

**Figure S10. Elevated expression of the DNA replication-associated gene signature is associated with tumor development.**

(A and B) The expression distribution of the DNA replication-associated gene signature across lung adenocarcinoma (A), osteosarcoma (B), and normal tissues from the TNMplot dataset. The DNA replication-associated gene signatures, including MCM3, MCM4, MCM5, MCM6, MCM7, RPA1, RPA1, PCNA, LIG1, RFC5, and FEN1, exhibit heightened expression in lung adenocarcinoma (A) and osteosarcoma (B) compared with normal tissues. The p-value is calculated using the Mann-Whitney test.

**Table S1. Spearman correlation analysis of KDM6A with MCM3, MCM4, MCM5, MCM6, MCM7, RPA1, RPA1, PCNA, LIG1, RFC5, and FEN1 in The Cancer Genome Atlas (TCGA), PanCancer Atlas.**
The significantly positive co-expression of KDM6A with DNA replication-associated gene signatures (with the positive Spearman correlation) is indicated in bold.

| Cancer type | Co-expression of KDM6A with DNA replication-associated gene signatures | | | | | | | | | | Database/ Case number |
|---|---|---|---|---|---|---|---|---|---|---|---|
| | MCM3 | MCM4 | MCM5 | MCM6 | MCM7 | RPA1 | PCNA | LIG1 | RFC5 | FEN1 | |
| Uterine Corpus Endometrial Carcinoma | **Spearman= 0.01; *P*= 0.0180** | **Spearman= 0.41; *P*= 7.90e-23** | Spearman= -0.06; *P*= 0.0150 | **Spearman= 0.16; *P*= 2.285e-4** | Spearman= 0.01; *P*= 0.865 | **Spearman= 0.43; *P*= 1.82e-25** | **Spearman= 0.11; *P*= 0.0150** | Spearman= -0.04; *P*= 0.409 | **Spearman= 0.20; *P*= 5.880e-6** | Spearman= 0.01; *P*=0.884 | TCGA, PanCancer Atlas/ n= 529 |
| Cervical Squamous Cell Carcinoma | Spearman= 0.06; *P*= 0.280 | **Spearman= 0.37; *P*= 3.96e-11** | Spearman= 0.02; *P*= 0.689 | **Spearman= 0.18; *P*= 2.223e-3** | Spearman= -0.03; *P*= 0.665 | **Spearman= 0.33; *P*= 7.17e-9** | **Spearman= 0.10; *P*= 0.0901** | **Spearman= 0.17; *P*= 2.926e-3** | **Spearman= 0.23; *P*= 6.308e-5** | Spearman= -0.09; *P*= 0.118 | TCGA, PanCancer Atlas/ n= 297 |
| Skin Cutaneous Melanoma | Spearman= 0.01; *P*= 0.831 | **Spearman= 0.13; *P*= 6.657e-3** | Spearman= -0.25; *P*= 1.18e-7 | **Spearman= 0.14; *P*= 3.497e-3** | Spearman= -0.12; *P*= 0.0114 | **Spearman= 0.15; *P*= 1.098e-3** | Spearman= -0.03; *P*= 0.577 | Spearman= -0.28; *P*= 3.26e-9 | **Spearman= 0.06; *P*= 0.220** | Spearman= -0.16; *P*= 7.026e-4 | TCGA, PanCancer Atlas/ n= 448 |
| Sarcoma | **Spearman= 0.24; *P*= 1.111e-4** | **Spearman= 0.41; *P*= 1.04e-11** | Spearman= 0.05; *P*= 0.393 | **Spearman= 0.31; *P*= 3.29e-7** | **Spearman= 0.21; *P*= 9.708e-4** | **Spearman= 0.40; *P*= 3.61e-11** | **Spearman= 0.16; *P*= 0.0102** | **Spearman= 0.27; *P*= 1.450e-5** | **Spearman= 0.35; *P*= 8.13e-9** | **Spearman= 0.14; *P*= 0.0313** | TCGA, PanCancer Atlas/ n= 255 |
| Lung Squamous Cell Carcinoma | Spearman= 0.02; *P*= 0.610 | Spearman= 0.04; *P*= 0.334 | Spearman= 0.04; *P*= 0.327 | **Spearman= 0.11; *P*= 0.0178** | Spearman= 0.01; *P*= 0.776 | **Spearman= 0.10; *P*= 0.0332** | Spearman= -0.04 *P*= 0.342 | **Spearman= 0.16; *P*= 2.849e-4** | Spearman= -0.03; *P*= 0.512 | Spearman= 0.00; *P*= 0.975 | TCGA, PanCancer Atlas)/ n= 487 |

#### Supplemental Table S2

| Antibodies | Source | Identifier |
|---|---|---|
| Rabbit polyclonal anti-UTX | Sigma-Aldrich | Cat# ABE409 |
| Rabbit monoclonal anti-ATM (D2E2) | Cell Signaling Technology | Cat# 2873 |
| Rabbit monoclonal anti-phospho-ATM (Ser1981) | Abcam | Cat# ab81292 |
| Mouse monoclonal anti-ATR (C-1) | Santa Cruz Biotechnology | Cat# sc-515173 |
| Rabbit monoclonal anti-phospho-ATR (Thr1989) | Cell Signaling Technology | Cat# 30632 |
| Mouse monoclonal anti-CHK1 (2G1D5) | Cell Signaling Technology | Cat# 2360 |
| Rabbit monoclonal anti-phospho-CHK1 (Ser345) | Cell Signaling Technology | Cat# 2348 |
| Mouse monoclonal anti-CHK2 (1C12) | Cell Signaling Technology | Cat# 3440 |
| Rabbit polyclonal anti-phospho-CHK2 (Thr68) | Cell Signaling Technology | Cat# 2661 |
| Rabbit monoclonal anti-phospho-H3Ser10 (H3S10ph) | GeneTex | Cat# GTX61067 |
| Mouse monoclonal anti-β-actin antibody, HRP conjugate | GenScript | Cat# A00730 |
| Rabbit polyclonal anti-GAPDH | Proteintech | Cat# 10494-1-AP |
| Goat anti-Rabbit IgG secondary antibody, HRP conjugate | Cell Signaling Technology | Cat# 7074 |
| Goat anti-Mouse IgG secondary antibody, HRP conjugate | Thermo Fisher Scientific | Cat# 31430 |
| Rat monoclonal anti-GFP (GF090R) | Nacalai USA | Cat# 04404-84 |
| Mouse monoclonal anti-PARP1 (B-10) | Santa Cruz Biotechnology | Cat# sc-74470 |
| Mouse monoclonal anti-phospho-Histone H2A.X (Ser139), clone JBW301 | Sigma-Aldrich | Cat# 05636 |
| Alexa Fluor 488 Goat Anti-Mouse IgG (H+L) | Thermo Fisher Scientific | Cat# A-11029 |
| Alexa Fluor 594 Goat Anti-Rabbit IgG (H+L) | Thermo Fisher Scientific | Cat# A-11037 |

#### Supplemental Table S3

| Oligonucleotides | Source | Identifier |
|---|---|---|
| **UTX (RT-qPCR)**<br>Forward- TACAAATCCGAACAACCCTG<br>Reverse- TGAGGAGGCCTGGTACTGT | This paper | |
| **MCM3 (RT-qPCR)**<br>Forward- TGGCCTCCATTGATGCTACC<br>Reverse- GGACGACTTTGGGACGAACT | Yang Laboratory | https://pubmed.ncbi.nlm.nih.gov/33093917/ |
| **MCM4 (RT-qPCR)**<br>Forward- CAGCAGCAAATCCCATTGAGT<br>Reverse- TGTCATAGGCTTCGTCCTGAG | Xue Laboratory | https://pubmed.ncbi.nlm.nih.gov/34961841/ |
| **MCM5 (RT-qPCR)**<br>Forward- TCATCTCCAAGAGCATCGCC<br>Reverse- CCTCGGCGAGTAAGTCCATC | Yang Laboratory | https://pubmed.ncbi.nlm.nih.gov/33093917/ |
| **MCM5 (RT-qPCR)**<br>Forward- TCATCTCCAAGAGCATCGCC<br>Reverse- CCTCGGCGAGTAAGTCCATC | Yang Laboratory | https://pubmed.ncbi.nlm.nih.gov/33093917/ |
| **MCM6 (RT-qPCR)**<br>Forward- GACAACAGGAGAAGGGACCTCT<br>Reverse- GGACGCTTTACCACTGGTGTAG | Wu Laboratory | https://pubmed.ncbi.nlm.nih.gov/32964028/ |
| **MCM7 (RT-qPCR)**<br>Forward- GCCAAGTCTCAGCTCCTGTCAT<br>Reverse- CCTCTAAGGTCAGTTCTCCACTC | Wu Laboratory | https://pubmed.ncbi.nlm.nih.gov/32964028/ |
| **RPA1 (RT-qPCR)**<br>Forward- GGGGATACAAACATAAAGCCCA<br>Reverse- CGATAACGCGGCGGACTATT | Jing Laboratory | https://pubmed.ncbi.nlm.nih.gov/34992431/ |
| **RPA3 (RT-qPCR)**<br>Forward- TTCGTAGGGAGGCTGGAAAA<br>Reverse- CCTTGGCGGTTACTCTTCCAA | Takami Laboratory | https://pubmed.ncbi.nlm.nih.gov/16919414/ |
| **RFC5 (RT-qPCR)**<br>Forward- GAAGCAGACGCCATGACTCAG<br>Reverse- GACCGAACCGAAACCTCGT | Bae Laboratory | https://pubmed.ncbi.nlm.nih.gov/33692216/ |
| **FEN1 (RT-qPCR)**<br>Forward- AGGGAGAGCGAGCTTAGGAC<br>Reverse- GAGGAGGGATGACTGGCCTT | This paper | |
| **LIG1 (RT-qPCR)**<br>Forward- TGGGAAGTACCCGGACATCA<br>Reverse- GCTTCGGTGTCCAGGATGAA | This paper | |
| **PCNA (RT-qPCR)**<br>Forward- TCTGAGGGCTTCGACACCTA<br>Reverse- CATTGCCGGCGCATTTTAGTA | This paper | |
| **RPL37A (RT-qPCR)**<br>Forward- CGACATGGCCAA ACGTACCA<br>Reverse- CTTGGCGTGCTGGCTGATTT | This paper | |
| **PCNA_enhancer (ChIP-qPCR)**<br>FW TGCACTCATTCCGTAAACACT<br>REV ACATGTGATGCTATGTTTTAAAGAGGT | This paper | |
| **LIG1_enhancer (ChIP-qPCR)**<br>FW CAGGAAATGACGTTGGCAGT<br>REV CGCCCAGCCTGTATTTTGTT | This paper | |
| **MCM3_enhancer (ChIP-qPCR)**<br>FW TCCTTCCTTGACATTCATACCCC<br>REV ACCCCAGTGTGGTACTTACA | This paper | |
| **PCNA_promoter (ChIP-qPCR)**<br>FW CCCACTTCCAAGCCATGTGT<br>REV CCACGCTTCAGTCTGCTTTG | This paper | |

| | |
|---|---|
| **LIG1_promoter (ChIP-qPCR)**<br>FW CCTCTGGGAGACACCTCTCT<br>REV AATGCTTGGGGCCTATGCTT | This paper |
| **MCM3_promoter (ChIP-qPCR)**<br>FW TGCAGCCGAATTATGGAGTCT<br>REV TTTGAGGAGACTGTGGGGGA | This paper |
